# RNase H genes cause distinct impacts on RNA:DNA hybrid formation and mutagenesis genome-wide

**DOI:** 10.1101/2023.05.08.539860

**Authors:** Jeremy W. Schroeder, Rebecca L. Hurto, Justin R. Randall, Katherine J. Wozniak, Taylor A. Timko, Taylor M. Nye, Jue D. Wang, Peter L. Freddolino, Lyle A. Simmons

## Abstract

RNA:DNA hybrids such as R-loops affect genome integrity and DNA replication fork progression. The overall impacts of naturally occurring RNA:DNA hybrids on genome integrity, and the relative contributions of ribonucleases H to mitigating the negative effects of hybrids, remain unknown. Here, we investigate the contributions of RNases HII (RnhB) and HIII (RnhC) to hybrid removal, DNA replication, and mutagenesis genome-wide. Deletion of either *rnhB* or *rnhC* triggers RNA:DNA hybrid accumulation, but with distinct patterns of mutagenesis and hybrid accumulation. Across all cells, hybrids accumulate most strongly in non-coding RNAs and 5′-UTRs of coding sequences. For Δ*rnhB*, hybrids accumulate preferentially in untranslated regions and early in coding sequences. Hybrid accumulation is particularly sensitive to gene expression in Δ*rnhC*; in cells lacking RnhC, DNA replication is disrupted leading to transversions and structural variation. Our results resolve the outstanding question of how hybrids in native genomic contexts interact with replication to cause mutagenesis and shape genome organization.

## INTRODUCTION

RNA:DNA hybrids (referred to herein as RDHs) form through a variety of essential processes such as transcription and DNA replication, giving rise to several distinct forms of RDHs *in vivo*. RDHs include single ribonucleotide misincorporation errors by replicative DNA polymerases, primers for Okazaki fragments during lagging strand replication, and R-loops formed when a single strand of RNA anneals to complementary DNA, displacing a strand of DNA in the process [reviewed in (*1–3*)]. RDHs contribute to genome instability in several ways. First, they are more stably paired than double stranded DNA (dsDNA), making their removal by helicases difficult. Second, RNA is much more susceptible than DNA to breaks due to the reactive 2’-hydroxyl group on the ribose sugar. Third, R-loops can pose barriers to DNA replication (*4–8*). The relative contributions of these biological and chemical properties of RDHs to genome instability are unclear. However, what is clear is that in organisms ranging from bacteria to humans, RDHs promote the accumulation of mutations that threaten genome integrity, cause double-strand breaks, and result in activation of the innate immune response in human cells (*6, 9–14*).

Because RDHs can be costly to cells, mechanisms exist to remove them. For example, in bacteria there are three types of RNase H enzymes capable of digesting the RNA strand of an RDH: HI, HII, and HIII (*15*). Most bacteria, including *Escherichia coli*, encode RNases HI and HII. However, an important subset of bacteria, including the Gram-positive bacterium *Bacillus subtilis*, contains RNases HII and HIII (*16, 17*). All three types of RNase H enzymes can incise ribonucleotides covalently embedded in DNA (*18, 19*). In contrast, only RNases HI and HIII incise substrates lacking a covalent RNA-DNA junction, suggesting these enzymes cleave R-loops *in vivo* (*18, 19*).

The mechanisms through which RNase H enzymes maintain genome stability are multi-faceted. RNase HII may reduce mutagenesis by enabling relatively error-free replacement of ribonucleotides mis-inserted by DNA polymerases (*10, 20, 21*). In *E. coli*, RNase HI suppresses double-strand break formation at sites of co-directional conflicts between DNA replication and transcription (*22*). Head-on replication-transcription conflict is especially deleterious (*7, 22–26*), and RNase HIII may reduce R-loop accumulation and mutation rates in head-on genes, potentially by helping to avoid replication fork reversal at head-on conflict sites (*7, 27*). Assigning relative importance to the contributions and interplay of RDHs, replication-transcription conflict, gene expression level, R-loop stability, gene location, and sequence context to mutagenesis has been difficult, in part due to the large number of biological inputs, and in part owing to the current lack of a highly specific, reproducible method for detecting RDHs.

To understand the impact of RDHs such as R-loops on genome integrity, it is necessary to have a highly specific method for RDH detection (*28*), while also accounting for confounding biological variables such as gene expression and transcription direction relative to DNA replication. We therefore used a recently developed, high-accuracy method for RDH detection and measured genome-wide RDH levels, alongside measurement of gene expression via RNA-seq, in *B. subtilis* cells lacking either RNase HII (*rnhB*) or HIII (*rnhC*). To relate these findings to mutagenesis, we integrated our gene expression and RDH abundance measurements with mutation accumulation line data using a statistical model that accounted for the diverse biological inputs affecting genome stability. We show that in the absence of RnhC, R-loops accumulate in regions of high expression and cause severe genome instability due to replication stress at a specific locus with pervasive RDH presence near the chromosomal replication terminus. By contrast, in cells lacking RnhB, RDHs accumulate preferentially in UTRs and early in coding sequences. Our integrative approach reveals that R-loops, regardless of gene orientation, promote accumulation of insertions and deletions in coding sequences in wild type cells. However, in nearly all cases, the selective pressures exerted by RDH formation appear to have minimized the potential for problematic conflicts in naturally evolved genes. Our combined analyses provide a genome-wide assessment for how transcription, DNA replication, and R-loops contribute to mutation rate and replication fork perturbations throughout a bacterial chromosome.

## RESULTS

### Loss of RnhB or RnhC results in global changes in RNA:DNA hybrid distributions

We began by testing the ability of *B. subtilis* RnhB (HII) and RnhC (HIII) to incise an R-loop substrate *in vitro* **(Figure 1A-B).** RnhB was not active against the R-loop, while RnhC efficiently cleaved the RNA strand of the R-loop substrate **(Figure 1A-B and Figure S1)**. Given the differing specificities of RnhB and RnhC enzymes *in vitro*, we expected that loss of *rnhB* and/or *rnhC* would influence the genome-wide abundance and distribution of RDHs in distinct ways. We used an epitope-tagged hybrid-binding domain (HBD) from human RNase H1 (*29*) to isolate RDHs with four or more ribonucleotides from wild type (PY79), Δ*rnhB*, and Δ*rnhC* cells followed by high-throughput sequencing of the input and RDH-enriched DNA. We quantified genome-wide RDH enrichment using a tool developed for this work called Enricherator (see Methods for details). We have shown that our HBD pull-down approach provides substantial improvements in sensitivity and specificity of enrichment relative to antibody-based RDH pulldowns (to be submitted elsewhere).

**Figure 1.**
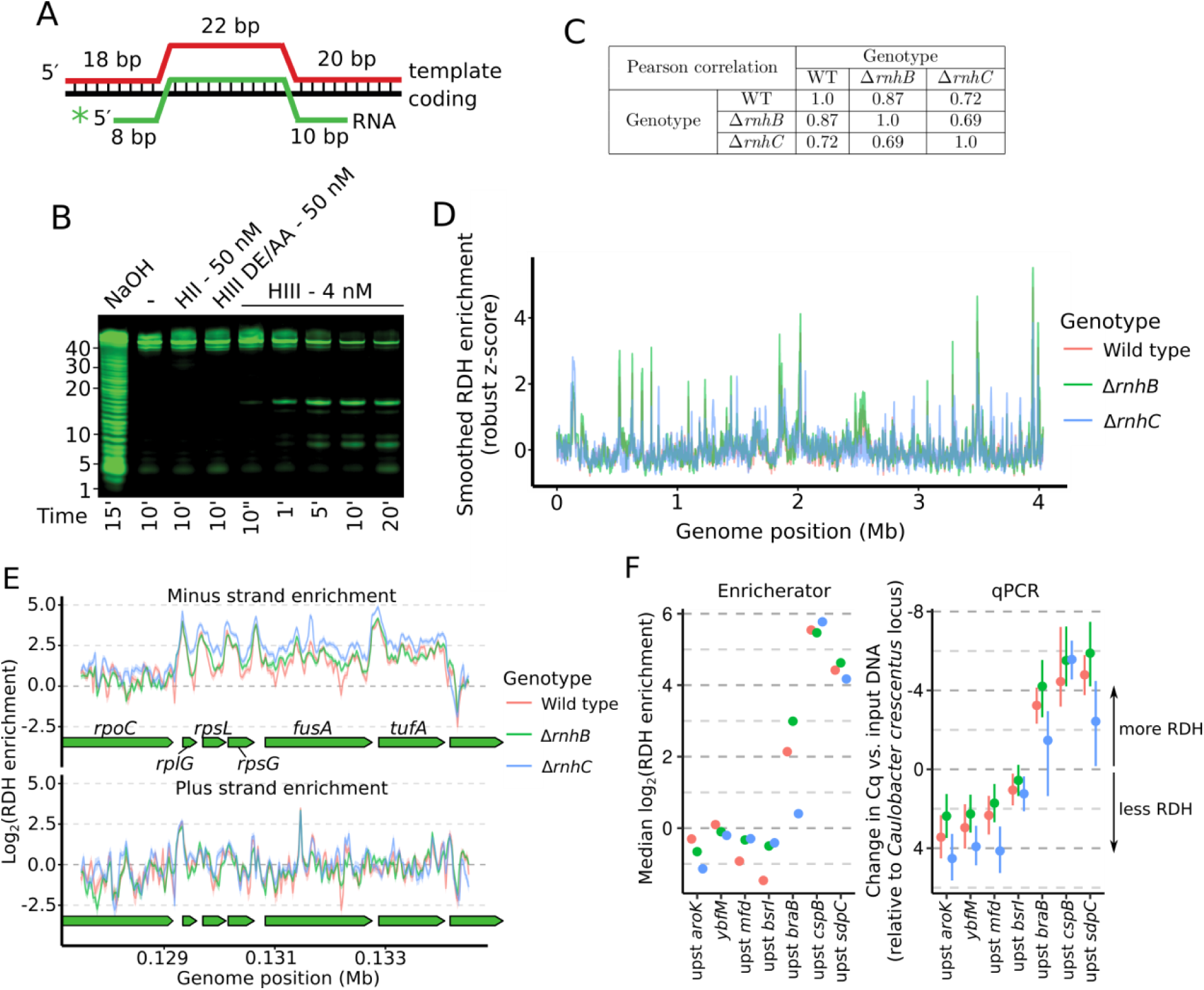
Loss of *rnhB* and/or *rnhC* causes widespread increases in RDH formation. **(A)** Diagram of the three oligonucleotides used to generate the R-loop substrate for *in vitro* cleavage reactions. (**B**) Urea-PAGE of the R-loop treated with 50 nM RnhB (HII), 50 nM catalytically inactive RnhC (HIII DE/AA), or 4 nM RnhC (HIII) at the indicated time points. (**C**) Pearson correlation coefficients for genome-wide comparison of RDH enrichment scores across genotypes. **(D)** Overview of RDH signal, smoothed using a 10 kb sliding median across the genome for each genotype. **(E)** Strand-specific RDH enrichment at a specific region of the genome. Lines represent the mean, and shaded intervals represent the 90% quantile interval of 500 samples from the approximate posterior distribution. Note that the shaded intervals are often narrow and are thus not always visible under the mean line. (**F**) Comparison of locus-specific enrichments inferred by Enrichterator (left) and as determined by qPCR (right). For the qPCR plot, points represent the mean of the samples from each posterior distribution and bars represent 95% highest posterior density intervals.

RDH enrichment scores varied over the genome similarly across genotypes, with Pearson correlation coefficients for any two genotypes between 0.69-0.87 **(Figure 1C-D)**. Wild type and Δ*rnhB* RDH enrichments were highly correlated indicating a similar distribution (Pearson correlation = 0.87), whereas Δ*rnhC* showed substantial differences in overall RDH density from Δ*rnhB* and WT. In general, highly transcribed regions were enriched for RDHs, especially in Δ*rnhC* cells **(Figure 1E)**. We sought to compare RDH accumulation across genotypes, and because our sequencing approach can only yield relative levels of enrichment within a given genotype, we performed qPCR of several genomic regions informed by our sequencing-based enrichments. To ensure we were comparing RDH enrichments to a known, unchanging reference locus, we spiked in an equal amount of *Caulobacter crescentus* cell pellet to each sample prior to lysing cells for input DNA and HBD pull-down procedures. HBD enrichment as determined by qPCR was defined in terms of ΔΔCq. Here, ΔΔCq is the difference in Cq between HBD pull-down vs. input DNA at a given *B. subtilis* locus, compared to the difference in Cq between HBD pull-down vs. input DNA at a single *C. crescentus* tRNA^Thr^ locus (reflecting a region with robust RDH formation). Comparing our genome-wide estimates from Enricherator and our targeted qPCR approach, differences in RDH pulldown efficiency across genotypes and loci were in excellent agreement between the two methods **(Figure 1F)**. Of note, most loci tested by qPCR displayed slightly lower RDH pulldown efficiencies for Δ*rnhC* cells than for wild type and Δ*rnhB* cells. Although based on the known biochemistry of RnhC and RnhB we initially expected the opposite to be the case (*19*), Δ*rnhC* cells are sensitive to several types of stress (see below and (*7*)), and we suggest that the absence of RnhC may globally reduce transcription due to the stress induced by loss of RnhC. Globally reduced transcription in Δ*rnhC* cells would in turn cause most loci to display lower RDHs per unit genome, such that only the loci with very high expression display higher RDHs in Δ*rnhC* cells.

### RNA:DNA hybrids are enriched at highly expressed genes, especially in cells lacking RnhC

In order to interpret the genome-wide patterns of RDH abundance observed in our experiments, we separately analyzed the distributions of RDH signals across several classes of genomic features (e.g., open reading frames, 3′-UTRs, etc.). Across all considered strains, ribosomal RNA loci had the highest levels of RDH accumulation, with 5′-UTRs having overall higher RDH accumulation than other features **(Figure 2A)**. Loss of *rnhC* further exacerbated RDH accumulation at rRNA loci.

**Figure 2.**
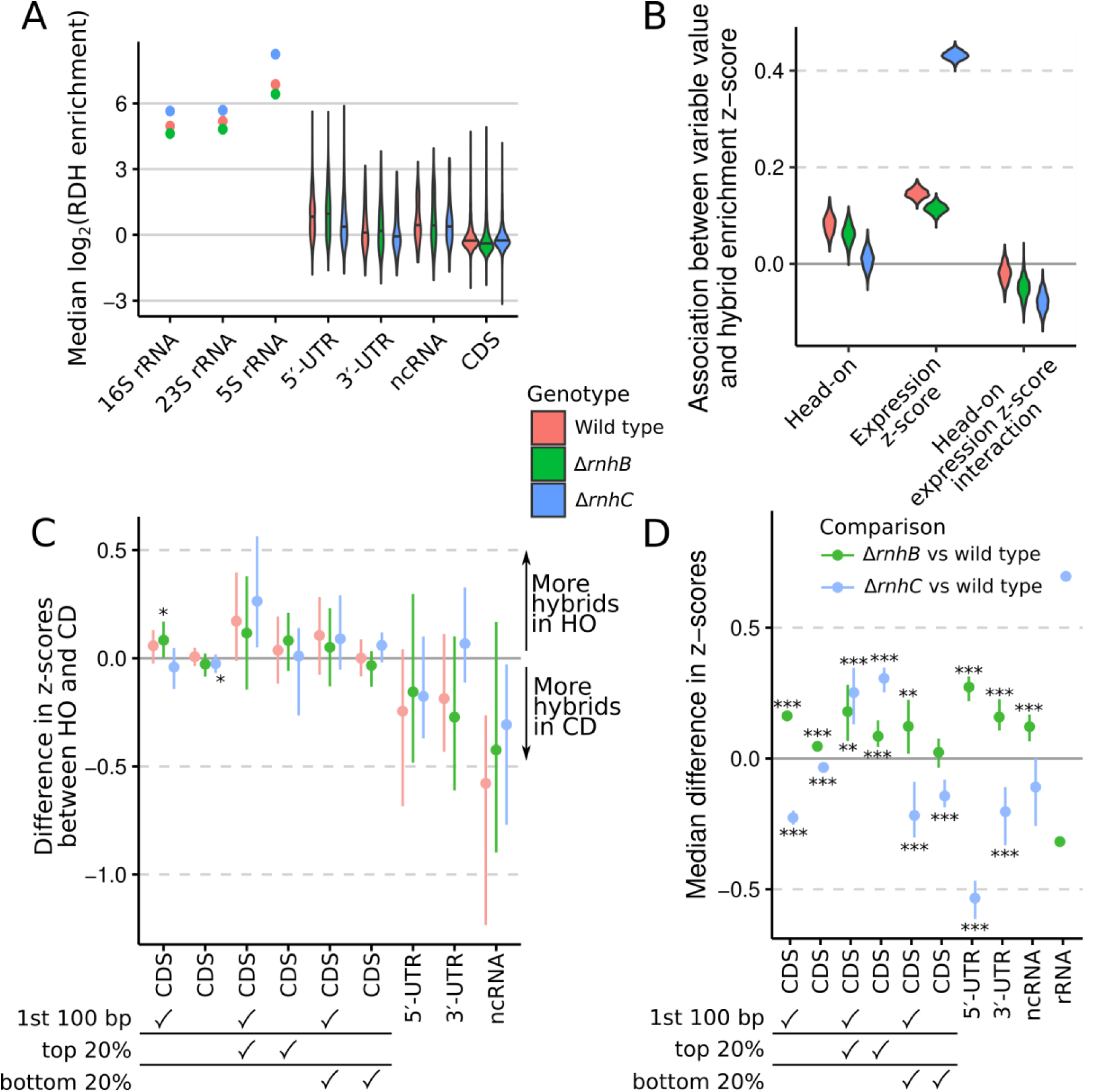
Loss of *rnhB* and/or *rnhC* causes changes to RDH signals for co-directional and head-on genes. RDH enrichment for each genomic feature was considered to be the median robust z-score for RDH enrichment within the feature. False discovery rate (FDR) indicators for panels C and D: *, 0.01 <= FDR < 0.05; **, 0.001 <= FDR < 0.01; ***, FDR < 0.001. Wilcoxon rank-sum tests were performed. **(A)** Distributions of RDH enrichment scores for each annotated feature of the type listed in the x-axis. For rRNA there is only one point per genotype because our custom reference genome contained a single consensus sequence for each rRNA, with the ten chromosomal copies masked. Therefore, for each rRNA a single enrichment score exists, whereas for other feature types we display the distributions of enrichment scores as violin plots. Horizontal lines across the violin plots represent the median value for each group. **(B)** Sampled posterior distributions are displayed for a model (see Supplemental Methods section SM5.5 for model details) regressing RDH enrichment z-score, aggregated by CDS, against gene orientation relative to DNA replication, gene expression z-score, and an interaction term between gene orientation and expression. The model was fit separately for each genotype. **(C)** The difference in head-on vs. codirectional feature RDH enrichment scores (points) is displayed with the 95% CI (bars). Check marks under the x-axis labels indicate whether filters for position within CDSs or gene expression were applied to the data for a given set of comparisons. **(D)** RDH enrichment z-scores were compared across genotypes, paired by genomic feature. The resulting median feature-wise difference in RDH enrichment is plotted (points) with the 95% CI (bars). Due to the low number of rRNA in our reference genome, we did not estimate confidence intervals or perform hypothesis tests for rRNA. Thus, the rRNA comparisons are plotted as points to indicate they represent only point estimates of the differences. For panels C and D, checkmarks in top 20% or bottom 20% denote that only genes in the top or bottom quintile expression were included.

Active expression of protein coding genes and the direction of gene transcription relative to that of DNA replication (head-on vs. codirectional) have been correlated with changes in R-loop enrichment (*7, 30*). We therefore expected that the most actively expressed genes should have increased RDH enrichment, and that the distribution of RDH scores for head-on and codirectional genes, and for the most actively expressed genes, may change in the absence of *rnhB* or *rnhC*. We fit a statistical model to regress RDH enrichment scores for CDSs against CDS expression, direction of transcription relative to DNA replication (head-on or codirectional), and the interaction between CDS orientation and expression. For all strains, increased CDS expression was associated with higher RDH enrichment scores, but this association was much more pronounced for Δ*rnhC* cells than for either wild type or Δ*rnhB* **(Figure 2B)**. For wild type and Δ*rnhB* cells, head-on orientation of a CDS was associated with a small increase in RDH enrichment, but increased gene expression in head-on genes had a lower association with RDH abundance **(Figure 2B)**, suggesting that existing highly expressed head-on genes in *B. subtilis* may have evolved to have lower R-loop accumulation than equivalent expression levels in co-directional genes would predict. Further we find no association between R-loops and loss of *rnhC* in head-on expressed genes in their native genomic contexts.

Next, we explored the association between gene orientation, expression, and RDH enrichment by separating genomic features into categories and calculating the difference in RDH enrichment z-scores between head-on and codirectional features. The first 100 bp of head-on CDSs, especially for highly expressed CDSs when *rnhB* was absent, displayed slightly higher RDH enrichment scores than codirectional CDSs **(Figure 2C)**. However, when the entire length of CDSs was considered, head-on and codirectional CDSs showed similar RDH accumulation, regardless of genotype. For non-coding RNA other than rRNA and tRNA, head-on transcription is associated with lower RDH enrichments **(Figure 2C)**.

Comparing RDH enrichment z-scores across genotype, we observed higher z-scores in Δ*rnhB* cells (relative to WT) across nearly all considered genomic features aside from ribosomal RNAs. Our finding is consistent with generally pervasive but less transcription-dependent RDHs in the absence of RnhB, since for highly expressed genes RDHs accumulate less in *ΔrnhB* than in *ΔrnhC* **(Figure 2B and D)**. Further underscoring the strong relationship between transcription and R-loop formation in Δ*rnhC* cells **(Figure 2B)**, RDH z-scores for Δ*rnhC* cells were higher than wild type in the 20% most highly expressed CDSs, but were otherwise lower than wild type when considering all CDSs, CDSs with low expression, and UTRs **(Figure 2D)**.

We conclude that while all cells accumulate RDHs strongly at highly expressed genes, RDH accumulation in the *ΔrnhC* genotype is more strongly affected by transcription, while RDH accumulation in *ΔrnhB* increases more evenly throughout the genome.

### Replication stress occurs at long head-on operons near the chromosomal terminus in Δ*rnhC cells*

As loss of *rnhB* or *rnhC* causes different patterns of RDH accumulation and increases in R-loop formation are associated with genomic instability, we expected that Δ*rnhB* and Δ*rnhC* cells would not only have higher mutation rates than wild type cells, but also that the types and frequencies of the mutations observed would differ in each genetic background. To understand the genome-wide contribution of RDHs to mutagenesis we performed mutation accumulation (MA) experiments using Δ*rnhC* cells, and compared mutagenesis in Δ*rnhC* to previously-published MA line results from wild type and Δ*rnhB* (*10, 31*).

For Δ*rnhC* cells, we completed 72 MA lines representing a total of ≈265,500 generations surpassed in the pooled lines. Among the genomic variations we detected in MA line data, we found that structural variants and transversions were particularly pronounced in Δ*rnhC* lines **(Figure 3A and Data File S2)**. The total number of structural variants detected in Δ*rnhC* was 17 compared to the 6 identified in wild type lines, resulting in a structural variation rate per generation of 6.4 x 10^-5^ for Δ*rnhC* compared to 2.2 x 10^-5^ for wild type (p = 0.015, one-tailed rate-ratio test; **Figure 3A)**. Structural variants clustered near the replication terminus in Δ*rnhC* lines **(Figure 3B-C)**. We reasoned that because mutations clustered near the terminus of Δ*rnhC* cells, DNA replication fork progression could be impaired in this region of the chromosome. To test this, we monitored DNA replication status genome-wide by performing whole-genome sequencing on DNA from exponentially growing cells. The slope of the genome sequencing coverage profiles served as a proxy for replication status in each strain, where a shallow slope was indicative of rapid replication fork progression, and a steep slope indicated replication proceeded with difficulty (*25*). For our genomic analysis of replication status we sequenced DNA from wild type, Δ*rnhC, lexA[G92D]*, and *rnhC::erm lexA[G92D]* cells. Cells harboring *lexA[G92D]* encode a non-cleavable LexA protein and are impaired for induction of the SOS response. We found that the genome-wide replication profiles were similar for wild type and *lexA[G92D]* cells, as indicated by similar slopes of genome sequencing coverage in each strain **(Figure 3B-C)**. However, in cells both lacking *rnhC* and impaired for activation of the SOS response (*rnhC::erm lexA[G92D]*), we found a steep drop in sequencing coverage near the terminus region of the genome where several head-on genes reside, including *yngEFG, yngIJ,* and *ppsABDE*, which is long, at about 27 kb **(Figure 3B-C)**. The drop in sequencing coverage indicates that DNA replication was substantially impaired as it approached the *ppsABDE* locus in the absence of *rnhC*. Replication stress in the *yng* and *pps* operons coincided with pervasive RDH increases in Δ*rnhC* relative to wild type cells in these head-on genes (**Figure 3D**), suggesting that DNA replication may be impaired at this locus due to pervasive R-loop accumulation.

**Figure 3.**
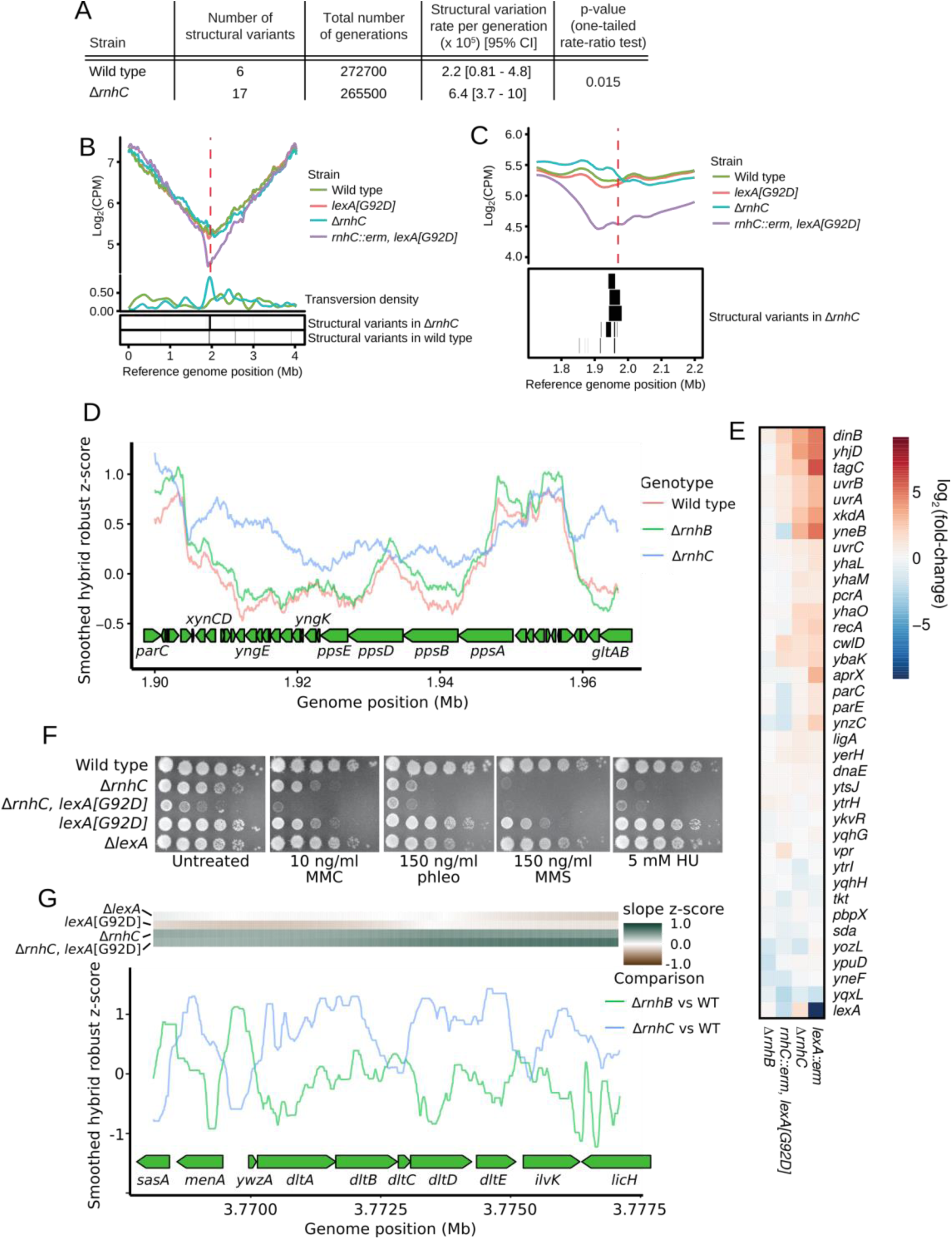
RnhC suppresses replication stress and genome instability. **(A)** Table showing structural variant data from wild type and Δ*rnhC* MA lines. **(B)** Log_2_ transformed sequencing read counts per million reads mapped (CPM) plotted versus genome position for wild type, *lexA[G92D]*, Δ*rnhC*, and *rnhC::erm lexA[G92D]*. Density of transversions is plotted (see Methods) below versus genome position for Δ*rnhC* and wild type. The red dashed line indicates the replication terminus. **(C)** Top - A view of the data plotted in B, zoomed in on the region near the replication terminus (red dashed line). Bottom - Structural variants detected in Δ*rnhC* mutation accumulation lines are plotted as boxes to indicate their genome position. **(D)** For each strain, RDH enrichment scores from Enricherator were converted to robust z-scores, smoothed using a 10kb rolling median, and plotted versus genome position near the *pps* operon. **(E)** Shown is a heatmap of gene expression for the SOS response in the indicated genetic backgrounds relative to wild type. **(F)** Spot titer assays of various RnhC and LexA-deficient strains serial diluted 10^-5^ and plated on LB with the chemical stress indicated in the figure. **(G)** Contrasts for each of Δ*rnhB* and Δ*rnhC* relative to wild type were prepared using Enricherator scores. Contrasts were converted to robust z-scores, smoothed using a 500 bp rolling median, and plotted at the *dlt* operon. The heatmap above the plot shows the z-score of the slope of the smoothed DNA sequencing coverage plotted in panel B. Dark green scores indicate decreased rate of DNA replication progression.

Inspection of the replication profiles suggests that the SOS response is induced in Δ*rnhC* cells. Indeed, using RNA-seq **(Figure 3E)** and single-cell reporters **(Figure S3A)**, we found that Δ*rnhC* cells were constitutively induced for the SOS response. RNA-seq results for the LexA regulon in Δ*rnhC* closely mirrored the Δ*lexA* (constitutive SOS) expression profile, whereas Δ*rnhB* cells did not show SOS induction **(Figure 3E)**. Further, we show that Δ*rnhC* cells were sensitive to hydroxyurea (HU) and a variety of DNA damaging agents when compared with wild type cells, sensitivities which were exacerbated by concomitant loss of SOS induction (**Figure 3F)**. Our findings that loss of *rnhC* leads to SOS induction and sensitivity to DNA damage support and extend previous findings underscoring the conclusion that Δ*rnhC* cells are severely compromised for genome integrity (*7, 19, 20*). We conclude that SOS induction is critical for survival and proliferation of cells lacking *rnhC*. Further, we suggest that in the absence of *rnhC*, increased mutagenesis near the terminus of replication is caused by replication stress due to defects in R-loop removal from the *yng* and *pps* operons in that region, followed by SOS induction.

In addition to replication problems near the *pps* operon, we detected a second site, the *dltABCDE* operon, where DNA replication proceeded slowly in cells lacking *rnhC* **(Figure 3G, top)**. The *dlt* operon is head-on and highly expressed, with the mean expression z-score of the genes in the operon ranging from 1.5 to 1.7 in wild type cells. Loss of *rnhC* caused increased and pervasive RDH accumulation in the *dlt* operon **(Figure 3G)**. Therefore, replication stress at the *dltABCDE* locus is likely the result of head-on replication-transcription conflict stabilized by R-loops. However, unlike the *ppsABDE* operon, which is 27 kb long and near the terminus, *dltABCDE* is less than 5 kb long, is distal to the terminus, and did not show any changes in mutation or structural variation rates. This comparison indicates that high expression, R-loop stabilization, and head-on orientation are not sufficient to increase rates of transversions and genomic rearrangements. We conclude that the elevation in R-loop formation at head-on genes negatively impacts DNA replication. To understand the impact of R-loops in head-on genes we tested whether there was a statistical association between mutagenesis, R-loops, and the direction of replication-transcription conflict (see below).

### RNA:DNA hybrids promote insertions and deletions genome-wide

We have demonstrated above that loss of *rnhC* causes widespread increases in RDH abundance, likely in the form of R-loops. We have also shown that SOS induction overcomes severe replication stress near the replication terminus in Δ*rnhC* cells. However, loss of RnhC has led to negligible mutation rate increases (1.3-fold) when mutagenesis is measured using mutation reporter genes (1.3-fold) (*7, 20*), and the genome-wide effects of loss of *rnhC* on mutagenesis are unknown. Therefore, to determine the genome-wide effect of Δ*rnhC* on mutagenesis we analyzed our MA line data for base-pair substitutions and small insertions and deletions (indels). The overall mutation rate for base-pair substitutions and indels in Δ*rnhC* was 1.6 x 10^-3^ **(Data File S2)** per generation, which is similar to that for Δ*rnhB* (*10*), and is roughly 50% higher than the wild type mutation rate of 1.1 x 10^-3^ (p < 0.001, rate-ratio test). Analysis of the types of mutations that occurred in Δ*rnhC* cells uncovered increased transition and transversion rates, with no strongly detectable difference in the accumulation of insertions or deletions genome-wide **(Figure 4A)**. The most striking mutagenic effect for base-pair substitutions was a 2-fold increase in transversions **(Figure 4A)**, a hallmark of the SOS response due to up-regulation of error-prone DNA polymerases, which are more likely to form transversions than replicative enzymes (*32*). Further, our results showing an increase in transversions are consistent with each of the following: first, the constitutive induction of the SOS response we observe in the absence of *rnhC* as described above, and second, the enrichment of transversions in the terminus region where replication forks become laggard and RNA:DNA hybrids are enriched **(Figure 3C)**.

**Figure 4.**
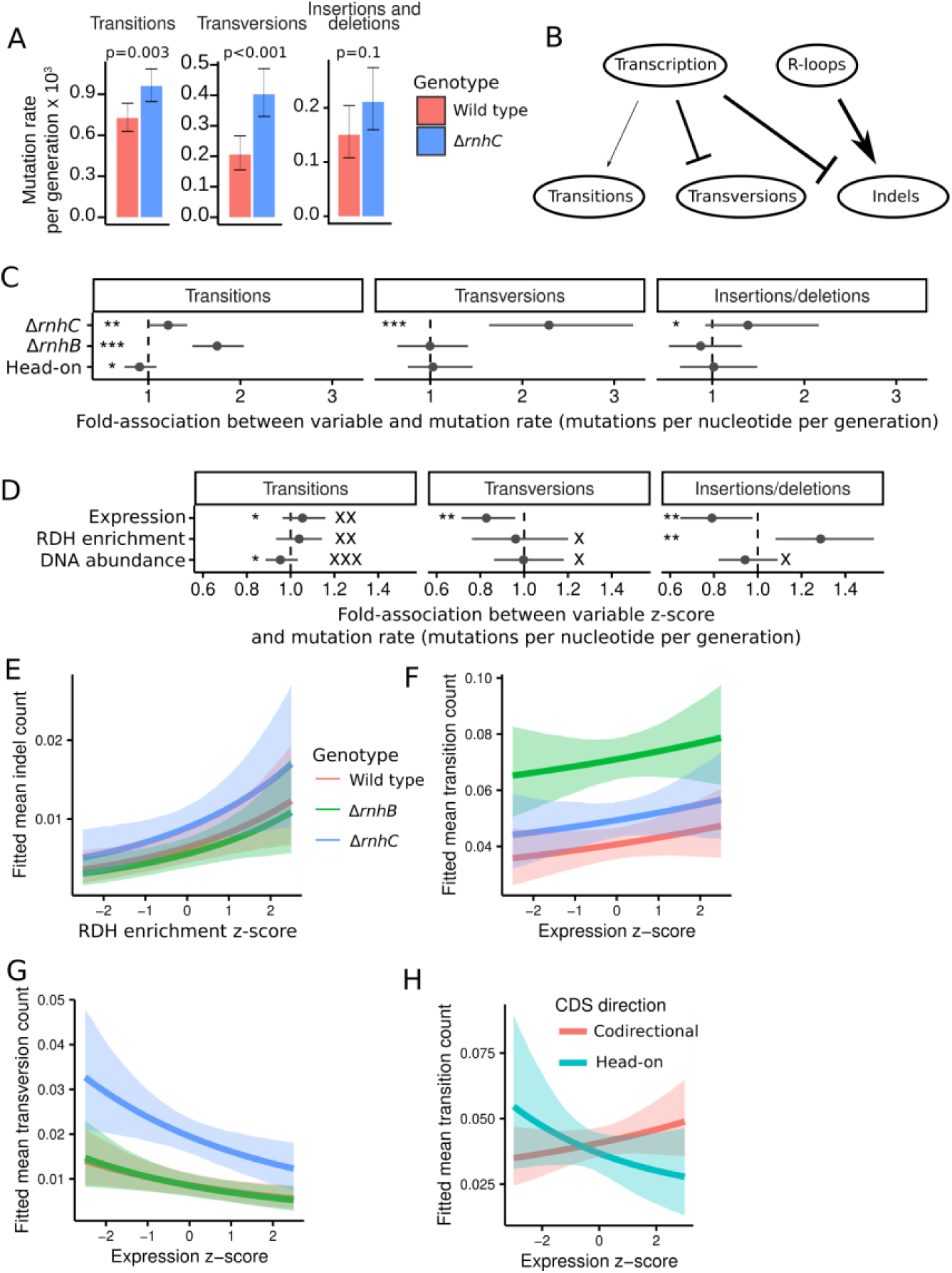
Effects of loss of RnhB and RnhC on mutation rate. **(A)** Bar plots indicating the genome-wide mutation rates derived from mutation accumulation line experiments for transitions, transversions, and insertions/deletions for wild type and Δ*rnhC* cells. Error bars represent 95% exact Poisson confidence intervals. **(B)** A schematic illustrating the relative contributions of transcription and R-loops to each type of mutation. Line weights are proportional to the median estimate of transcription or R-loops association with the indicated mutation type. Arrows indicate a positive association, and flat heads denote a negative association. **(C)** Inferred associations between the rate of transition, transversion, or indel occurrence in CDSs and the indicated model parameters for categorical variables. **(D)** Inferred associations between mutation rate in CDSs and the indicated model parameters for continuous variables. All associations for interaction terms are in Figure S4A. **(E)** Plot showing the mean fitted number of transitions in CDSs with varying gene expression and all other variables held at their mean. The shaded regions indicate the 90% quantile interval for the fitted values. **(F)** A plot showing the mean fitted number of transversions in CDSs with varying gene expression and all other variables held at their mean. **(G)** A plot showing the mean fitted number of indels in CDSs with varying RDH abundance and all other variables held at their mean. **(H)** A plot showing the relationship between mean fitted number of transitions in CDSs and gene expression and all other variables held at their mean, stratified by head-on or codirectional. Throughout the figure, *, **, and *** symbols indicate the strength of evidence for a parameter having a substantial association with mutation rate, with * indicating K is from [3,20), ** denoting K is from [20,150), and *** indicating K ≥ 150. Equivalently strong K_0_ values are shown with equal numbers of “X” symbols. Note that if a parameter can be very precisely estimated but is close to zero, it is possible for it to have both high K and high K_0_.

With the diverse types of data we collected, we fit a Bayesian statistical model to identify associations between mutation rate in CDSs and several variables, including RDH abundance, expression, and CDS direction relative to replication (head-on vs. co-directional). In contrast to the simple rejection of a null hypothesis provided by frequentist methods, our Bayesian approach enabled us to quantify the strength of evidence in favor of each parameter’s value being positive, negative, or nearly equal to zero. We summarize the strength of evidence for each hypothesis in terms of Bayes factors (K), for which a high K-value (> 150) indicates very strong evidence, K-values from 20-150 indicate strong evidence, and K-values from 3-20 indicate positive evidence (*33*). In the discussion below, K is used to indicate whether a given effect is substantial, whereas K_0_ tests the hypothesis that the effect of interest is near zero. Thus, a high K_0_ indicates a significant lack of any notable effect. Several aspects of mutagenesis in wild type, Δ*rnhB*, and Δ*rnhC* MA lines were revealed using this approach (see Supplementary Materials and Methods SM5 for method and statistical models, see **Data File S2** for a summary of the inferred parameter estimates). For example, the fit of our statistical model corroborates our conclusion that loss of *rnhC* increases transversion, transition, and indel rates (K > 150, K = 24.6, and K = 8.6, respectively) **(Figure 4A, 4C)**. In agreement with prior work (*10*), loss of *rnhB* increased CDS transition rate (K > 150) due to a previously described error prone ribonucleotide excision repair pathway (*10*) **(Figure 4C)**.

Increasing RDH abundance was associated with increased indel rate (K = 59.6) but did not affect transition or transversion rates (K_0_ = 68.0 and 4.1, respectively) **(Figure 4B and D).** Although it has been suggested that R-loops are particularly mutagenic in head-on genes, our data suggest that on average, across all native CDSs in the *B. subtilis* genome, associations between mutagenesis and RDH accumulation were independent of gene orientation relative to DNA replication **(Figure S4A and S4D)**. We interpret this to indicate that in contrast with engineered mutation reporters designed to detect mutations due to head-on conflict (*7, 24*), native genes that have evolved in a head-on genomic context have done so such that the influence of head-on transcription and R-loop formation on mutagenesis is minimized. Thus, engineered reporters can demonstrate the underlying molecular mechanisms driving the selective pressures on gene orientation, but in practice selective pressures have eliminated most potentially damaging conflicts in bacterial genomes.

Increased gene expression has also been associated with increased mutagenesis. We found that gene expression had a very small positive association with transition rate (K = 5.1, K_0_ = 46.6), whereas transversion and indel rates were lower as expression increased (K = 38.2 and 23.1, respectively) **(Figure 4B-C, F-G)**. Additionally, for head-on genes our data show the opposite association between transition rate and gene expression. Overall, we show that higher expression of head-on genes correlates with a decreased transition rate **(Figure 4G, Figure S4A** and **S4C,** and discussion). We conclude that loss of *rnhC* results in an increase in transition, transversion and indel mutations while loss of Δ*rnhB* yields an increase in transitions. Further, we conclude that R-loop-dependent mutagenesis in Δ*rnhC* is independent of gene orientation and is instead dependent on the extent to which RDHs accumulate and the length of a genomic locus over which they accumulate.

We demonstrated above that transcription and RDH accumulation are correlated **(Figure 2B)**, therefore we tested whether the correlation between gene expression and RDH accumulation affected our ability to separately infer the association between each variable and mutation rates by fitting two new statistical models. Each was identical to the model described above, with the exception that the first new model lacked any terms associated with gene expression, and the second lacked any terms associated with RDH enrichment. If either variable affected inferences about the other’s association with mutation rate, then this approach would have detected that relationship. However, the inferred associations between expression or HBD accumulation and mutation rate did not change appreciably when leaving the other variable out of a given model, and the predictive performance as measured by LOO-CV was not influenced by leaving either variable out of the model (see Supplemental Methods Section SM6 for details). Therefore, although expression is correlated with RDH abundance, each influences mutagenesis separately. Knowing both expression level and RDH abundance will provide a much better prediction of the gene’s mutation rate than knowing either parameter in isolation.

## DISCUSSION

RNA:DNA hybrids (RHDs) arise in many different forms and have been shown to affect replication fork progression, mutation rate, and cell survival (*7, 10, 20, 21, 34*). Here we investigate the consequences of loss of either *rnhB* or *rnhC in vivo* to RDH accumulation, replication fork progression, and genome-wide mutagenesis. The major finding of our work is that R-loops increase indel rates, whereas loss of *rnhC* leads to increased transversion rates due to R-loop-induced replication stress and induction of the SOS response. We are now able to propose models for the roles of RnhB and RnhC in RDH removal and their consequences for genome stability in bacteria.

Altogether, we suggest that the major benefits of R-loop digestion by RnhC are to limit replication stress and fork reversal **(Figures 3-4** and (*27*)), and to keep 3′-termini of R-loops from being used inappropriately as primers for DNA synthesis. RnhB can act on such inappropriate priming events if R-loops escape the activity of RnhC or if cells lack *rnhC* entirely (*19, 35*). Cells lacking *rnhB* showed an increase in RDH abundance in UTRs and in the first 100 bp of CDSs. However, we demonstrate that purified RnhB shows little activity on an R-loop lacking a covalent RNA:DNA junction *in vitro*, while RnhC shows robust activity. Therefore, we propose that increased RDH abundance at UTRs and early in CDSs in Δ*rnhB* cells represents the fraction of R-loops that escape RnhC action and are used to prime DNA synthesis at sites of replication fork blockage. Under normal conditions, these priming events generate a covalent RNA-DNA junction leading to resolution by RnhB. However, in the absence of *rnhB* the re-priming events are stabilized and detected in our pulldown assay. This model would also explain how RDH in *rnhB* cells show little dependence on transcription.

We attempted to gain additional insights into RDH formation by deleting both *rnhB* and *rnhC*. While it is possible to knock out both *rnhB* and *rnhC* in a single *B. subtilis* strain, Δ*rnhB*Δ*rnhC* cells have a slow growth rate (*20*). We have also found that each time we sequence the genome of a Δ*rnhB*Δ*rnhC* isolate, we identify compensatory mutations (*35*). This suggests that loss of both *rnhB* and *rnhC* together leads to an exceedingly high mutations rate which quickly leads to accumulation of suppressor mutations, causing us to abandon further work on a Δ*rnhB*Δ*rnhC* strain.

We show that Δ*rnhC* cells are exquisitely sensitive to DNA damage [**Figure 3F, Figure S3B** and (*19*)] and accumulate RDHs in both gene orientations in a transcription-dependent manner. We considered the possibility that growth inhibition of Δ*rnhC* in response to DNA damage was due to failed primer removal from Okazaki fragments, which was compounded by the addition of exogenous DNA damage. If this had been the case, then overexpression of RnhB or Pol I would have provided moderate rescue, as RnhB and Pol I are involved in ribonucleotide excision repair (RER) and Okazaki fragment processing in *B. subtilis* (*10, 20, 35*). However, neither *rnhB* nor Pol I (*polA*) overexpression rescued any sensitivity (**Figure S3B**), suggesting that the sensitivities were primarily due to deficiencies in R-loop resolution, an activity in which *B. subtilis* RnhC has been implicated [**Figure 1A-B** and (*7*)]. We find it likely that the SOS response is nearly saturated in Δ*rnhC* cells, which would cause both their sensitivity to DNA damaging agents and the synergistic sensitivity of Δ*rnhC lexA[G92D]* cells to those same agents.

In addition to DNA damaging agents, Δ*rnhC* cells are sensitive to osmotic stress, cold sensitivity, lysozyme, oxidative stress, and hydroxyurea (*7, 19, 35*). One model that has been proposed to explain the phenotype is that loss of *rnhC* may impair induction of head-on stress-induced genes (*7*). Another possible explanation arising from the work presented here is that persistent, genome-wide R-loop formation in all genes, independent of orientation **(Figure 2B-C)**, causes DNA replication stress that is further exacerbated by an exogenously applied stress condition.

While prior work in *B. subtilis* has suggested that *rnhC* overexpression may decrease mutagenesis in head-on reversion reporter genes (*7*), studies in yeast using head-on and co-directional reporter genes have shown that replication-transcription conflicts caused by R-loops are independent of gene orientation (*36*). Our approach has the advantage of querying genome-wide mutagenesis and RDHs in native gene contexts. Understanding the effect in native gene context is important because native genes have already evolved under selective pressure from effects of gene position and orientation on mutagenic potential. Our work shows that RDHs mainly increase indel rates in coding sequences and promote transversions in Δ*rnhC* cells where replication forks encounter R-loop impediments. An important feature of our finding is that the transversion mutations identified in *B. subtilis* are enriched in the *pps* operon located near the replication terminus.

The consensus view of the association of gene expression with mutagenesis is that as gene expression increases so does mutagenesis (*37–41*), and while our inference corroborates prior evidence that increased transcription increases the transition mutation rate of a gene, we found that the effect is likely to be very small in the genes natively found in *B. subtilis*. The stronger effects of transcription on mutagenesis are the negative associations between expression and transversion and indel rates. We assert this negative effect of transcription on transversion and indel rates is likely due to transcription-coupled nucleotide excision repair acting on the types of DNA damage that would otherwise cause transversions and indels [reviewed in (*42, 43*)].

The types of mutations caused by, and the evolutionary consequences of, head-on replication-transcription conflict and conflict between DNA replication and R-loops in head-on genes is a subject of ongoing discussion (*7, 24, 31, 36, 44, 45*). Bacterial genes are enriched in the co-directional orientation because of negative selection against head-on conflicts which disrupt DNA replication and cause mutagenesis (*46*). Therefore, head-on genes tend to be those that are either more tolerant to mutation or produce fewer conflicts [reviewed in (*46, 47*)], and the few native head-on genes that do produce significant conflicts will likely be purified from the genome over time. By contrast, synthetic mutation reporter genes have not been subject to the evolutionary pressures shaping the genome (*7, 24*) and are able to provide insights into how mutagenesis is affected by extreme, engineered conflicts that select for specific mutations. Engineered gene conflict reporters are thus best used to illustrate the mechanisms underlying the selective pressures that have acted to purify or minimize head-on conflicts in native genes. Although the results presented here and elsewhere (*48*) do not demonstrate a general association between gene direction and mutagenesis, we have identified one native locus, *ppsABDE*, that produces results similar to those from engineered mutation reporters. Without *rnhC*, *ppsABDE* accumulates transversions and structural variants.

The *pps* locus also exhibits significant replication stress in Δ*rnhC* cells, and the increased transversion rate can be attributed to induction of the SOS response, which we show is required for replication to proceed through *ppsABDE* in Δ*rnhC* cells. Because SOS induction includes the up-regulation of Y-family polymerases that are prone to causing transversion mutations when replicating DNA [reviewed in (*32*)], increased transversions and detrimental structural variants near the terminus suggest that SOS induction and Y-family polymerases are involved in DNA replication at this chromosomal location in Δ*rnhC* cells. Therefore, when specific conditions are met, including head-on orientation, stable R-loop formation, considerable length, SOS induction, and proximity to the replication terminus, then one native locus in *B. subtilis* shows increased mutagenesis in the head-on orientation. The mutations arising in the *pps* operon are unlikely to provide an adaptive benefit as they are often large deletions.

Taken together, our whole-genome data allow reconciliation and mechanistic explanation of a wide range of factors related to the interplay of RDH formation, genome organization/gene orientation, and mutagenesis that have been observed in different experimental systems. Global analysis of mutagenic profiles demonstrates that regardless of their origin, RDHs are associated with increased insertion/deletion mutations. While loss of function of either *rnhB* or *rnhC* causes increases in RDH accumulation and overall mutation rates, the differing effects of loss of the two genes arise due to resolution of different types of RDHs, resulting in different types of mutations (transitions vs. transversions and indels) and different biases in where RDHs accumulate. For naturally occurring genes under baseline physiological conditions, a head-on orientation relative to transcription is not mutagenic, but loss of *rnhC* function unmasks the mutagenic potential of very specific head-on oriented genes when R-loop formation is increased. In general, it appears that the *B. subtilis* genome has reached a point of equilibrium with respect to gene orientation, in which regularly expressed genes in the head-on orientation have undergone selective pressure to minimize R-loop formation and consequently mutagenesis. These findings explain the discrepancy between mutagenesis rates observed in long, highly expressed artificial reporters vs. native genes.

## MATERIALS AND METHODS

### Bacteriology

All strains used in this study are described in **Data File S1** in the supporting material. Cells were grown in S7_50_ defined minimal medium (*49*) or LB medium at 30°C. For antibiotics, cells with *recA-gfp* were selected for using 100 µg/mL spectinomycin, and 5 µg/ml erythromycin. Further details are given in Supplementary Materials and Methods SM1.

### Spot titer assays

Assays were performed essentially as described in (*19*). Briefly, a single colony of the indicated strain was used to inoculate 3 ml of LB media and grown to an OD600 between 1 and 1.5. Cultures were then normalized to an OD600 = 1 in a 0.85% saline solution and serial diluted to 10^-5^. A total volume of 5 μl was spotted for each dilution on LB or LB agar containing the indicated concentrations of MMC, MMS, Phleomycin, HU and or IPTG. Plates were then incubated overnight at 30°C and imaged the following morning.

### Fluorescence microscopy

RecA-GFP and TagC-GFP strains were imaged as described previously (*50, 51*). See Supplementary Materials and Methods SM2 for further information.

### Mutation accumulation lines

Procedures were performed as described (*10, 31*). Previously published wild type (*B. subtilis* PY79) and Δ*rnhB* MA line data were compiled (*10, 31*). Structural variation in MA lines were detected using the software “breakdancer” (*52*). Structural variants detected within 100 base pairs of the intended genetic alteration for an MA line were removed from further analysis. For instance, *rnhB* has the start and end positions 1640489 and 1641256, respectively, so structural variants detected by breakdancer within the location from genomic coordinates 1640389 to 1641356 were removed from this study. Upon inspection of the structural variants in our MA lines, we noticed that a Δ*rnhC* line contained a deletion of the 3′ end of the *mutS* gene. Because *mutS* is required for DNA mismatch repair, the mutation rate and spectrum would be strongly affected (*31*) by this structural variation in *mutS*. Therefore, we removed the line from this study.

### Replication assessment using genome sequencing

Three biological replicates each of strains JWS208, JWS266, JWS207, and JWS268 (see **Data File S1** for strain descriptions) were grown to mid-exponential phase in LB at 37°C. Cells were harvested via centrifugation, pellets were washed with PBS, and genomic DNA was purified. Genomic DNA sequencing libraries were prepared by the University of Michigan Sequencing Core, and single-end sequencing was performed on an Illumina Hiseq 2500 instrument. The number of fragments aligning at each position of the genome was determined. Sequencing counts were converted to log2(counts-per-million), and the mean log2(counts-per-million) was calculated across biological replicates for each strain. Mean log2(counts-per-million) were subsequently smoothed for visualization using the loess function in the statistical software R with the span argument set to 0.03.

### Plotting transversion density

To calculate and plot the transversion densities used in Figure 3 we used the geom_line function from the R package ggplot2, with the arguments stat=”density” and adjust=0.2.

### Genome-wide profiling of RNA:DNA hybrids

Details of the preparation of epitope-tagged hybrid binding domains, cell growth for pulldown experiments, sequencing library preparation, and subsequent data analysis, are given in Supplementary Materials and Methods SM5.

### Cell lysis and nucleic acid extraction

RDH pull-down experiments were performed by first extracting total nucleic acid from snap-frozen cells of the appropriate genotypes. Equivalent *Caulobacter crescentus* spike-in pellets (1 per sample) were resuspended in 1x low salt lysis buffer (660 μL/pellet; 10 mM Tris HCl, pH 8.0; 50 mM NaCl), transferred to a single *B. subtilis* pellet, resuspended via pipetting, and sonicated four times for 5 sec at 25% amplitude with a 15 sec pause between pulses (Branson digital sonifier). Then 1/10 volume of 10% SDS and 10 μL of proteinase K (50 mg/mL; EO0491; Thermo Scientific) were added to each sample, mixed, then incubated 55°C for 5 min. Then 1/10 volume of 3M sodium acetate and 500 μL of phenol/chloroform/isoamyl-alcohol was mixed in each tube. Tubes were incubated with mixing (600 rpm) for 6 min at 65°C . After sitting on ice for 5 min, the layers were separated via centrifugation (10 min; 16,000 r.c.f.; 4°C). The water layer was transferred to a new tube containing 500 μL of chloroform, vortexed, then spun 5 min, 16,000 x g at 4°C. The aqueous layer was transferred to a new 2 mL microfuge tube and ≥ 2.5 volumes of ice-cold, 100% ethanol added. Tubes were incubated -20°C for 60 min. Nucleic acids were pelleted by centrifugation (15 min; 16,000 x g; 4°C), supernatant removed, the pellet washed once with 80% ethanol, and then air dried until ethanol was absent. Nucleic acids were dissolved in 10 mM Tris pH 7.5. This solution was stored at -80°C until the following step.

### Digestion of nucleic acids for RNA:DNA extraction from cells

To each tube/sample was added 20 μL of 10X MNase buffer (New England Biolabs Inc.; Ipswich MA, USA), 0.5 μL of 1M MgCl2, 2 μL RNase inhibitor (M0314; New England Biolabs Inc.; Ipswich MA, USA) and 2 μL MNase (M0247; New England Biolabs Inc.; Ipswich MA, USA) were mixed in and tubes promptly placed on ice. After 20 min, 10 μL of 500 mM EGTA was added to each sample. The concentration of nucleic acid for each sample was determined by measurement of A_260_ (1 to 100 dilution). Aliquots of 500 µg were made of each sample. Then 1/10,000 of the sample, 50 ng of previously folded RNA-DNA hair-pin spike-in (P2918; Data File S1) was added to each sample. For most of these samples, 1X PBST was added to achieve a 200 µL volume and mixed by vortexing. Then 0.5 µL of 1 MgCl_2_ and 2µL RNase inhibitor (NEB) were added to each sample and mixed by vortexing. A 20μL aliquot per sample was placed in a new tube as an input sample then stored at -80° C. The remaining digested nucleic acids were used directly for HBD mediated pull-down as described below.

### HBD mediated pull-down of extracted and digested RNA:DNA hybrids

Freshly thawed on ice, purified HA-tagged HBD protein was added to the remaining nucleic acid digestion from each sample and mixed by repeated inversion. The HBD proteins were allowed to bind to the digested nucleic acids for 20 min at room temperature. Magnetic anti-HA beads (25 μL/tube; Pierce) were washed twice briefly with 1x PBST (137 mM NaCl, 12 mM phosphate, 2.7 mM KCl, pH 7.4, 0.2% Tween-20), liquid removed, and replaced with the remaining nucleic acid digestion. Tubes were incubated at room temperature on the undulating shaker for 20 min. Unbound material was removed. Beads were washed with 200 μL of 1x PBST and with 200 μL of 1x PBST+850mM NaCl_2_, then resuspended in 100 μL of 1x PBST and moved to a new tube. After the 1x PBST was removed, beads were resuspended in 50 μL of 1x TE and incubated at 95°C for 5min. Samples were returned to a magnetic stand, allowed to clear, and liquid transferred to DNA LoBind tubes (Eppendorf). Samples were stored at -80°C.

### Next-generation sequencing and data analysis

Illumina sequencing libraries were prepared from the extracted and input samples (see Supplementary Materials and Methods SM5) and sequenced on an Illumina NextSeq 500 instrument using 39x38 bp paired end reads. Data were subsequently processed using an in-house pipeline in which read alignments were processed to yield extracted:input ratios at each position on the chromosome, followed by normalization to robust z-scores. Details are given in Supplementary Materials and Methods SM5.

### Statistical modeling of mutagenesis

A detailed description of the model used to infer parameter estimates for mutation rate can be found in Supplementary Materials and Methods, Section SM6.

### *Purification of RnhB and RnhC for* in vitro *characterization*

All RNase H proteins were purified and assayed as described previously (*10*). The R-loop digestion buffer used 4 nM RnhC, 50 nM RnhB or 50 nM RnhC catalytic mutant in the following reaction buffer 20 µL of RNase H buffer (10 mM Tris-HCl pH 8, 50 mM NaCl, 1 mM MgCl_2_ or 10 µM MnCl_2_). For *in vivo* relevant metal concentrations both 1 mM MgCl_2_ and 10 µM MnCl_2_ were used as described previously (*19*).

### R-loop substrate formation and RNase H assay

The R-loop substrate was based on a previously published R-loop substrate (*53*). The R-loop was assembled by first annealing 10 µM oJR336 and oJR335 in a 50 µl solution of annealing buffer (10 mM Tris-HCl pH 8, 50 mM NaCl) by heating to 80°C and allowed to cool to 25°C. Then oJR332 was added to 10 µM and the solution was heated to 40°C and again allowed to cool to 25°C. The product was then visualized by native-PAGE in an 8% gel the R-loop substrate was gel extracted and diluted to 0.5 µM final concentration. Gel extraction and verification of the substrate are described in detail in the supporting information.

### Data availability

New sequencing data used in this study are available at the Sequence Read Archive (SRA) under BioProject accession number PRJNA962887.

## ACKNOWLEDGEMENTS

We thank Dr. Eileen Brandes for constructing the *lexA[G92D]* allele, Will Hirst for constructing strain WGH35 (Δ*rnhC, tagC::tagC-GFP*), Dr. Nicole Mott for initial method development work and for purification and characterization of the HBD-HA fusion protein, and Dr. Christine Ziegler for assistance with protein purification. J.R.R. was supported in part by a pre-doctoral fellowship from the Rackham Graduate School at the University of Michigan and NIH Cellular Biotechnology Training Grant (T32 GM008353). J.W.S. was supported in part by a pre-doctoral fellowship NIH T32 GM007544 as part of the Genetics Training Program. T.M.N. and K.J.W. were supported by National Science Foundation predoctoral fellowships (#DGE1256260). In addition, K.J.W. was supported in part by the NIH Cellular Biotechnology Training Program (T32 GM008353). Research in the laboratory of L.A.S. was supported by NIH grant R35 GM131772. Research in the laboratory of P.L.F. was supported by NIH grant R35 GM128637.

## Notes

### Competing Interest Statement

The authors have declared no competing interest.

