## Supplemental Methods, Table and Figures for "RNase H genes cause distinct impacts on RNA:DNA hybrid formation and mutagenesis genome-wide"

[1]Department of Biological Chemistry, University of Michigan Medical School, Ann Arbor, MI 48109. [2]Department of Molecular, Cellular, and Developmental Biology, University of Michigan, Ann Arbor, MI 48109. [3]Department of Bacteriology, University of Wisconsin - Madison, Madison, WI 53706. [4]Department of Molecular Microbiology and Center for Women's Infectious Disease Research, Washington University School of Medicine, Saint Louis, MO 63110-1093, USA.

‡To whom correspondence should be addressed:

Lyle A. Simmons, Department of Molecular, Cellular, and Developmental Biology, University of Michigan, Ann Arbor, Michigan 48109-1055, United States. Phone: (734) 647-2016, Fax: (734) 615-6337.

Peter L. Freddolino, Department of Biological Chemistry, University of Michigan, Ann Arbor, Michigan 48103, United States. Phone (734) 647-5839, Fax: (734) 763-4581.

Running Title: R-loops compromise genome integrity
Keywords: R-loops, mutagenesis, RNase HIII, *Bacillus subtilis*, mutation accumulation lines, SOS

### Supplementary Materials and Methods

*SM1: Bacteriology and cloning*

Strain JRR38 was created by transforming PY79 with pJR71 and plating on LB agar supplemented with 100 µg/ml spectinomycin. JWS266 was created by transforming PY79 with pEB17 and plating for *erm*$^R$ with growth at 37 °C. Colonies were then inoculated in 5 ml liquid LB and grown at room temperature, back-diluting every 12 hours for two days. 30 µl of this culture was then used to inoculate 5 ml of liquid LB and grown to an $OD_{600}$=1.1 and then diluted $10^{-6}$ in 0.85% saline. 100 µl of this dilution was plated on LB agar and grown overnight at 37 °C. Colonies were then screened for loss or *erm*$^R$ and genotyped for *lexA[G92D]*. JWS267 and JWS 268 were created by ordering a *B. subtilis* 168 *lox-erm*$^R$*-lox* knockout of *lexA* and *rnhC* loci respectively from the *Bacillus* Genetic Stock Center. Genomic DNA from these strains was purified and used to transform chemically competent PY79 (JWS267) or JWS266 (JWS268) and plated for *erm*$^R$. JRR67 and WGH35 were created by transforming genomic DNA from LAS40 and LAS366 respectively into chemically competent JWS224 and plating for *spec*$^R$.

Plasmid pEB16 was created by amplifying the *lexA* locus from PY79 gDNA using oEB64 and oEB65 and annealing it to the pMiniMAD2 vector via sequence and ligation independent cloning (SLIC). pEB17 was generated by an overlapping PCR reaction of pEB17 using oEB58 and oEB59 as primers. pJR71 was generated by amplification of the *polA* locus from PY79 gDNA using oJR260 and oJR261and cloning it into the pDR110 vector via Gibson assembly.

*SM2: Fluorescence Microscopy*

The use and imaging of RecA-GFP and TagC-GFP have been described in detail previously (*50*, *51*). Briefly, strain LAS40 (*recA-gfp*), TAT4 (Δ*rnhC*, *recA-gfp*), LAS366 (*tagC::tagC-GFP)*, or WGH35 (Δ*rnhC, tagC::tagC-GFP)* were grown in defined $S7_{50}$ minimal medium supplemented with 2% glucose at 30℃ from a starting $OD_{600}$ of 0.05 to an $OD_{600}$ of 0.4. Cells were placed on 1% agarose pads made in 1 x Spizizen's salts. Cells were imaged with an Olympus BX61 microscope at 1000 x magnification.

*SM3: In vitro assessment of RnhB/C activity*

*SM3.1: Purification of RnhB and RnhC*

The purification of RnhB and RnhC was performed as described (*10*). Briefly, each gene was cloned into pE-SUMO and overexpressed with a SUMO tag in *E. coli* $BL21_{DE3}$ cells at an OD~1 at 37$^{o}$C in LB with 200 µM IPTG. Cells were further grown at 37°C for 3 hours. Cells were lysed via sonication and cell debris was pelleted via centrifugation. The cell lysate was passed over a $Ni^{2+}$-agarose column, washed, and eluted with 400 mM imidazole. The eluate was then dialyzed into a buffer containing SUMO protease and left overnight at 4°C (50 mM Tris HCl, pH 8.0, 50 mM NaCl and 2 mM beta-mercaptoethanol) The SUMO cleaved protein was then passed back over a $Ni^{2+}$-agarose column and the flow through was applied to an Q-anion exchange column (GE: 17115301) and eluted with a NaCl gradient of 50 to 1000 mM. Fractions were analyzed via SDS-PAGE and the fractions containing the desired protein were pooled, exchanged into protein storage buffer (50 mM Tris HCl, pH 8.0, 100 mM NaCl, 25% glycerol), and flash frozen in liquid nitrogen.

*SM3.2: Polyacrylamide gel extraction of the R-loop substrate.*

The R-loop containing band was identified using a LI-COR Odyssey imager and was extracted. The band was extracted by the “crush and soak” method with incubating in 100 µl of elution buffer (300 mM magnesium acetate and 1 mM EDTA pH 8.0) rotating overnight at 4°C. The tube was centrifuged for 1 minute at 13,000 rpm and the supernatant containing the R-loop was removed. The yield was quantified by the intensity of the signal on the LI-COR Odyssey imager.

*SM3.3: Mung bean nuclease digestion*

Mung bean nuclease (MBN) was purchased from NEB (#M0250S). A concentration of 50 nM oJR336, oJR332, or R-loop was digested with MBN for 1 minute in a 10 µl reaction and quenched with stop buffer (95% formamide, 5 mM EDTA, 0.01% bromophenol blue). 0.3 M NaOH was added to oJR336 and used as a ladder. Reactions were heated to 100°C for 2 minutes and immediately snap cooled in an ice water bath. 20% Urea-PAGE was performed on all samples and digested products were imaged using a LI-COR Odyssey imager.

*SM3.4: RNase H digestion*

RNase H digestions were performed in 20 µL of RNase H buffer (10 mM Tris-HCl pH 8, 50 mM NaCl, 1 mM $MgCl_2$, 10 µM $MnCl_2$). Each reaction contained 0.5 µM of the R-loop substrate and either 50 nM RnhB, 50 nM RnhC DE/AA, or 4 nM RnhC. Reactions were performed for the indicated times and reactions were quenched with a stop buffer (95% formamide, 5 mM EDTA, 0.01% bromophenol blue). Reactions were heated to 100°C for 2 minutes and immediately snap cooled using an ice bath. 20% Urea-PAGE was performed on all samples and digested products were imaged using a LI-COR Odyssey imager. 0.3 M NaOH digestion of the R-loop was used as a ladder and to map the sites of cleavage.

*SM4: RNA-seq experiments and data analysis*

Three independent cultures each of JWS105, JWS207, JWS267, and JWS268 were grown to mid-exponential phase in LB at 37°C. Cells were immediately harvested by centrifugation after addition of one volume of ice-cold methanol. Total RNA was purified according to the manufacturer's instructions using the RiboPure RNA purification kit (Life Technologies). Ribosomal RNA depletion was performed by the University of Michigan DNA Sequencing Core, using the Ribo-Zero Magnetic Kit, Bacteria (Illumina), in accordance with the manufacturer's instructions. cDNA preparation, sequencing library preparation, and sequencing of 50-base single-end reads on an Illumina Hi-seq instrument were performed by the University of Michigan DNA Sequencing Core. Reads were aligned to the *B. subtilis* PY79 reference genome (CP006881.1) using bwa, version 0.7.8-r455. RNA-seq data from wild type *B. subtilis* PY79 were taken from previously published data (*31*) (SRA accession SRP067020).

Analysis of differential expression was performed using the statistical software R. Overlaps between coding sequences and alignments were counted using the summarizeOverlaps function from the GenomicAlignments package (*54*), with the arguments ignore.strand set to TRUE and mode set to IntersectionNotEmpty. Voom normalization was performed using the packages edgeR and limma (*55*, *56*) . P-values were adjusted for multiple testing by the method of Benjamini and Hochberg (*57*). The probability of differential expression was calculated by applying the logistic transform to the log-odds of differential expression for each gene returned by the topTable function in the package limma.

*SM5: Genome-wide profiling of DNA:RNA hybrids* (*expanded from main text*)

*SM5.1: Cell growth for RNA:DNA hybrid pulldown experiments.*

*B. subtilis* strains were streaked on LB plates and grown overnight at 37°C. In the morning, 3 mL of LB was used to resuspend cells from the plate and the optical density at 600 nm (OD) was determined. A larger LB medium culture was then inoculated to achieve an OD ~0.009. The larger culture was grown at 37°C in a baffle bottom flask until reaching OD ~0.6. Cells harvested for the RNA:DNA hybrid extraction and recovery were placed on ice and then split into 50 mL portions. Cells were pelleted via centrifugation for 4 min at 5500 x g at 4°C. Media was decanted, followed by the resuspension of cells in 10 mL of ice-cold, 1X PBS. Cells were pelleted via centrifugation for 4 min at 5500 x g at 4°C. Buffer (1X PBS) was decanted, cells resuspended in residual buffer and then transferred to one pre-chilled 2 mL microfuge tube per aliquot. Cells were collected via centrifugation (4°C; 2 min; 16,000 x g ). After buffer removal via micropipette, cells were flash frozen in an ethanol/dry ice bath. Cell pellet samples were stored at -80°C until processed further.

*Caulobacter crescentus* (LS101, obtained from Prof. Lucy Shapiro) were grown overnight in PYE medium at 30° C and then used to inoculate 160 mL PYE medium to achieve a density of 0.003 $A_{600}$. Grown at 30°C in a baffle bottom flask, cells were ultimately harvested at $A_{600}$ = 0.618, when two 50 mL aliquots were subjected to centrifugation (5500 rcf, 4° C, 4 min). After decanting, the two pellets were resuspended in 0.5mL of ice-cold 1x PBS, combined, and brought to 2mL total. The 50 uL aliquots were snap frozen for use as spike-in with *B. subtilis* samples which underwent HBD pull-down. The exact same batch of aliquots were used as spike-ins for all experiments described here.

*SM5.2: Purification of tagged HBD for genome-wide pulldown experiments*

Overnight culture of BL21 Gold cells containing a plasmid for expressing HA/His tagged HBD (HBD--MFYAVRRGRKTGVFLTWNECRAQVDRFPAARFKKFATEDEAWAFVRKSASPAGAGEQKLISEEDLGAGAYPYDVPDYAGSGHHHHHH) from an arabinose inducible promoter (NM003; Data File S1) were diluted 1:100 in induction medium [2% tryptone; 0.5% yeast extract; 0.6% glycerol; 0.2% arabinose; 0.082% glucose; 0.72% $Na_2HPO_4$; 0.28% $NaH_2PO_4$; 1x MOPS micronutrient mix (*58*) ] with chloramphenicol (30 μg/mL). Culture was grown at 37°C for ~3.5 hr then at 30°C until cells reached 4 to 5 $OD_{600}$ (16-48 hrs). Cells were washed with a Tris based buffer (50 mM Tris pH 7.5, 300 mM KCl), and transferred to 50 mL conical tubes (2 per 500 mL culture). The cells were collected via centrifugation (5500 r.c.f., 4° C, 20 min), decanted, weighed, stored -80° C).

NM003 cell pellets (28g) were thawed then resuspended in 280 mL of buffer (50 mM Tris pH 7.5, 300 mM KCl) by inversion and shaking at 4° C. Cells were sonicated for a total of 6 min (5s on, 25s off) at level 7 (Sonicator 3000, Misonix). Solution was split between 6 - 50 mL conical tubes, then insoluble material pelleted via centrifugation (12500 r.c.f., 4° C, 30 min). Soluble material was decanted and placed on ice. Elution buffer (15.8 mL of 50 mM Tris pH 7.5, 300 mM KCl, 250 mM imidazole) was added to the soluble material to achieve a final concentration of 12.5 mM imidazole. Nickel column was pre-washed with 50 mL of equilibrium buffer (50 mM Tris pH 7.5, 300 mM KCl, 12.5 mM imidazole). Half of the buffered soluble material was loaded onto the column, followed by a gravity driven gradient formed from equilibrium buffer (50 mM Tris pH 7.5, 300 mM KCl, 12.5 mM imidazole) and elution buffer (50 mM Tris pH 7.5, 300 mM KCl, 250 mM imidazole). This pre-wash, sample loading, and elution were repeated with the second half of the soluble material. Fractions containing significant amounts of the HBD protein were pooled, then concentrated using 4 - 3000 kD molecular weight cut off centrifugation columns (3200 r.c.f., 4° C, 10 min per spin). All nickel purified material was pooled, buffered with the Q-sepharose equilibration buffer [50 mM Tris pH 8, 1 mM EDTA pH 8, 0.1 mM 1,4-dithiothreitol (DTT)] to dilute the KCl to 30 mM. The 50 mL Q-sepharose column was pre-washed with 250 mL equilibration buffer [50 mM Tris pH 8, 1 mM EDTA pH 8, 0.1 mM 1,4-dithiothreitol (DTT)]. The buffered nickel purified material was loaded, followed by a gravity driven gradient formed from Q-sepharose equilibrium buffer (50 mM Tris pH 8, 1 mM EDTA pH 8, 0.1 mM DTT) and Q-sepharose elution buffer (50 mM Tris pH 8, 1 mM EDTA pH 8, 0.1 mM DTT, 0.5 M KCl). Fractions containing significant amounts of the HBD protein were pooled, then concentrated using 3000 kD molecular weight cut off centrifugation columns (3200 r.c.f., 4° C, 10 min per spin) to a final volume of 3 mL. The 10X concentrated storage buffer (529.4µL of 200 mM HEPES pH 7, 1 M KCl, 5 mM EDTA, 10 mM DTT) and 60% glycerol (1.765 mL) were added to the concentrated Q-sepharose purified protein. An additional 6.25 mL of 1x storage buffer with 20% glycerol was added to the buffered Q-sepharose purified protein, and gently mixed. The protein was further concentrated in the same 3000 kD molecular weight cut off centrifugation columns (3200 r.c.f., 4° C, 10 min per spin), aliquoted, and stored -80° C.

*SM5.3: Preparation and sequencing library production for HBD pull-down*

Each sample was brought to 50 µL volume using 1x TE. Then 2 µL of RNaseA (10 mg/mL) was added and mixed into each sample. After a 30 min incubation at 37° C, Oligo Clean & Concentrator kit (Zymo Research) kit was used to clean all samples. Briefly, to each sample, add 100 µL binding buffer and 400 µL 100% ethanol and vortex. Load onto columns. Centrifuge columns (12k rcf, 30s) and then empty collection tube. Add 750 µL of wash buffer to columns, centrifuge columns (12k rcf, 30s), and then empty collection tube. Dry column by centrifugation (12k rcf, 30s). Transfer column to DNA low-bind tube, add 20 µL of TEe to membrane, wait 30s, centrifuge columns (12k rcf, 60s), and then retain sample. Heat samples to 95° C for 3 min, ice, then use QuantiFluor ssDNA System (Promega) to determine ng/µL. Calculate volume (and dilutions required) to add 10 ng of DNA (and water) to achieve 14.5 µL for ssDNA sequencing preparation.

Sequencing preparation was performed as described by (*59*) in the STAR Methods section starting at "MNase-SSP" using 10 ng of DNA. The exact sequences of the "PCR_AMP_P7" and "PCR_AMP_P5" used for our samples are detailed in Data File S1. These libraries were cleaned up using 1.8x AxyPrep Mag PCR beads and quantified using QuantiFluor dsDNA System (Promega). Prepared libraries were stored at -20°C.

*SM5.4: Computational processing of RNA:DNA pull-down sequencing data*

The raw sequencing data was preprocessed by the removal of PCR duplicate reads, adapter sequences, and low-quality bases and reads. To facilitate removal of PCR duplicate reads we included 6bp unique molecular identifiers (UMIs) at the 5′ end of our R2 reads. We removed duplicate reads using pardre (command line flags: -l 8 -c 15 ) (*60*). We next trimmed the UMIs from the R2 reads and then trimmed adapters from de-duplicated reads using CutAdapt (command line flags: --quality-base=33 -a AGATCGGAAGAGC -A AGATCGGAAGAGC --mask-adapter -n 3 --match-read-wildcards). We then removed low-quality bases and reads with TrimmomaticPE (Command line flags: -phred33 LEADING:3 TRAILING:3 SLIDINGWINDOW:4:15 MINLEN:10). All samples were aligned to an in-house version of the

PY79 genome in which repetitive regions of the genome were hard masked and consensus 5S, 16S, and 23S rRNA were included as separate records within the reference fasta file. Alignment was performed using Bowtie version 2 (Command line flags: -q --end-to-end -X 2000 --very-sensitive --no-unal --phred33 –dovetail –fr). All code used to create our custom reference genome and its associated annotations can be found in the following github repository (https://github.com/jwschroeder3/hbd_enrichment_code.git)

The above preprocessing and alignment were performed separately for each of two sample types for each biological replicate: a sample taken just prior to the pull-down (input) and the material resulting from the HBD pull-down sample (extract). There are two biological replicates per genotype (PY79, Δ*rnhB,* Δ*rnhC)*. After alignment, the count of fragments aligning to each position of the genome at 5bp resolution was determined. To infer strand-specific pull-down enrichment at each genome position, we used a newly developed Bayesian analysis method implemented in Stan (*61*) which we call "Enricherator" (code can be downloaded at https://github.com/jwschroeder3/enricherator.git).

*SM5.5: Bayesian inference of genome-wide RNA:DNA hybrid enrichment scores*

Enricherator uses a variational Bayesian method to fit a statistical model with a negative-binomial likelihood to genome-wide input and extracted DNA sequencing coverage over input DNA sequencing coverage. Here, "extracted" can mean any type of experiment in which the user would like to determine enrichment of their target of interest. In the present work "extracted" refers to our HBD pull-down sequencing data, but it could be ChIP-seq, etc. for others' use cases. Enricherator produces point estimates and quantile intervals (by default, a 90% highest posterior density interval estimate) for input DNA abundance and extract enrichment by default and can optionally produce similar point estimates and quantile intervals for contrasts between strands or for contrasts between genotypes of interest, providing direct inference of the biological differences between samples.

Facets of Enricherator that make it preferable to most current methods of calculating genome-wide enrichment of extracted DNA signal over input DNA signal are: 1) The use of negative binomial statistics explicitly handles count data appropriately without the need for pseudocounts. 2) Enricherator applies a regularized horseshoe prior to enrichment scores, thus explicitly handling the Bayesian analog of multiple hypothesis testing. This shrinkage prior allows users to directly use the enrichment estimates arising from Enricherator runs without the need to apply shrinkage post-hoc. 3) Enricherator uses information from nearby genomic positions to inform its estimate of enrichment at any given genome position. It achieves this by using a smoothing kernel to mix nearby enrichment estimates. 4) The precision term to the negative-binomial likelihood is fit separately for input sequencing counts and extract (HBD pull-down, ChIP-seq, etc.) sequencing counts, providing maximal model flexibility to appropriately handle biological data. 5) Enricherator uses variational inference to approximate a posterior distribution of enrichment and input DNA scores. Use of variational inference allows the model to fit on the order of 7-8 hours of wall time for bacterial genomes, whereas sampling from the posterior distribution using conventional Monte Carlo methods would take a prohibitively long time. A description of the statistical model follows, and the full source Stan code and helper R scripts can be found at the Enricherator github repository referenced above.

We define subscripts in our model specification as follows:

| subscript symbol | definition | variable type |
|---|---|---|
| $p$ | genome position | ordinal |
| $g$ | genotype | categorical |

| | | |
|---|---|---|
| *t* | sample type (input or pull-down) | indicator |
| *s* | strand | indicator |
| *j* | sequencing library | categorical |

We model the number of fragments arising, $\boldsymbol{k}$, as a negative binomial, with expectation $\boldsymbol{\mu}$ and a separate global precision term, $\boldsymbol{\phi}$, for each sample type:

$$k_{p,g,t,s,j} \sim \mathrm{NB}(\mu_{p,g,t,s}, \phi_t)$$

We calculate the expected number of fragments, $\boldsymbol{\mu}$, for each combination of position, genotype, sample type, and strand as follows:

$$\ln(\mu_{p,g,t,s}) = \alpha_{p,g,s} + t \times \beta_{p,g,s} + c_j$$

$$\beta_{p,g,s} = \gamma_g + \delta_{p,g,s}$$

$$\gamma \sim \mathrm{T}(3, 0, 5)$$

where $\boldsymbol{\alpha}$ represents the the expected input DNA coverage for the given combination of position, genotype, and strand, $\boldsymbol{\beta}$ represents the enrichment of HBD pull-down signal for the given combination of position, genotype, and strand, *t* is an indicator variable for whether the sample is of the "extract" type, and $c_j$ is the ln(library size) for sequencing library *j*, centered on the mean ln(library size) across all sequencing libraries (see section "Library size calculation" below for details on calculation of $c_j$). Therefore, the term $c_j$ is an exposure term which performs library size normalization. The symbol $\boldsymbol{\gamma}$ represents a vector with a single element for each genotype analyzed. Each element of $\boldsymbol{\gamma}$ is a global offset for the enrichment scores within a given genotype. We set a weak prior on $\boldsymbol{\gamma}$ using a Student-T distribution (represented by the function "T" above) with 3 degrees of freedom, mean of 0, and standard deviation of 5. Above, $\boldsymbol{\delta}$ represents the deviation from the global extract enrichment in genotype *g* at genome position *p* for each strand *s*. Our use of a global genotype-specific enrichment offset $\boldsymbol{\gamma}$ and a position-wise vector of offsets for each genotype $\boldsymbol{\delta}$ enabled our use of a shrinkage prior on $\boldsymbol{\delta}$, described in greater detail below. The vectors $\boldsymbol{\alpha}$ and $\boldsymbol{\delta}$ are at the genomic resolution of the original input count data, and the values in these vectors are not inferred parameters. Rather, values in $\boldsymbol{\alpha}$ and $\boldsymbol{\delta}$ are calculated from inferred values in lower-dimensional, i.e., lower-resolution, vectors $\boldsymbol{\alpha'}$ and $\boldsymbol{\delta'}$ (see section "Accounting for covariance between nearby genome positions" below for details on calculating $\boldsymbol{\alpha}$ and $\boldsymbol{\delta}$).

**Library size calculation.**

The centered ln(library size) for each sequencing library was calculated by first taking a trimmed mean fragment count for each sequencing library, inspired by a similar approach implemented in the popular RNA-seq analysis R package edgeR (*62*). We trimmed the bottom 2.5% and the top 2.5% coverage data from the observed position-wise counts, $\boldsymbol{k}$, in each library and calculated the mean of the remaining counts. Below, $n$ represents the number of genome positions analyzed at 5 bp resolution after the lower and upper quantiles described above have been excluded. The trimmed mean ln(library size) for library $j$, $LLS_j$ was next centered on the mean of the vector of $\boldsymbol{LLS}$ across all libraries to calculate the exposure term, $c_j$, for each sequencing library.

$$LLS_j = \ln\left(\frac{\sum_{2.5\%ile}^{97.5\%ile} k_{p,j}}{n}\right)$$

$$c_j = LLS_j - \text{mean}(\boldsymbol{LLS})$$

**Accounting for covariance between nearby genome positions**

Sequencing data is generated in such a way that the number of observed fragments (*i.e.*, the "counts") at nearby genomic positions have a high degree of mutual information, in that knowing the coverage *k* at position *i* will give high confidence that coverage at position *i+1* will be closer to *k* than the coverage at a randomly-selected genome position. This covariance between coverage at nearby positions exists primarily due to the fact that entire fragments of DNA sequence are aligned to a reference genome, such that a single alignment will increase counts at several nearby genome positions simultaneously. We wrote Enricherator to account for covariance between nearby genome positions in both input sequencing libraries and extract sequencing libraries. In this section we use the terms *IFL* and *EFL* as variables to indicate "mean input fragment length" and "mean extract fragment length", respectively. *IFL* is used to define the degree to which estimates of input sequencing coverage are smoothed, i.e., the strength of covariance between nearby positions, and *EFL* is used to define the degree to which estimates of enrichment are smoothed. Higher values for each will yield greater smoothing. Details of how *IFL* and *EFL* are used to account for covariance between nearby genomic positions' input coverage and enrichment estimates, respectively, are below.

Consider that for a given sample type, genotype, and strand, $\boldsymbol{\alpha}$ and $\boldsymbol{\delta}$ are each a vector of genome positions of size (genome length)/resolution. Let us call mean input fragment length *IFL* and mean extract fragment length *EFL*. We define two new vectors, $\boldsymbol{\alpha}'$ and $\boldsymbol{\delta}'$, each element of which represents a fitted value at the following resolutions: *IFL*/2 and *EFL*/2. These reduced-dimensionality vectors of fitted values are next used to calculate vectors $\boldsymbol{\alpha}$ and $\boldsymbol{\delta}$ by deconvolving nearby values of $\boldsymbol{\alpha}'$ and $\boldsymbol{\delta}'$, respectively, using an exponential kernel. The decay rate of the exponential kernel is *IFL*/2 for deconvolving $\boldsymbol{\alpha}'$ to $\boldsymbol{\alpha}$ and *EFL*/2 for deconvolving $\boldsymbol{\delta}'$ to $\boldsymbol{\delta}$ **(Figure S2)**.

The above description is defined in the equations that follow, with $W_{\alpha}$ representing the sparse matrix of deconvolution weights for calculating $\boldsymbol{\alpha}$ from $\boldsymbol{\alpha}'$, and $W_{\delta}$ representing the sparse matrix of deconvolution weights for calculating $\boldsymbol{\delta}$ from $\boldsymbol{\delta'}$. Matrix $W_{\boldsymbol{\alpha}}$ has shape (size($\boldsymbol{\alpha}$), size($\boldsymbol{\alpha'}$)) and $W_{\boldsymbol{\delta}}$ has shape (size($\boldsymbol{\delta}$), size($\boldsymbol{\delta'}$)), and for each matrix, row values are normalized to sum to 1.0. The lower-dimension vector $\boldsymbol{\alpha'}$ was assigned a normal prior with mean 0 and standard deviation 4 (annotated as "N(0,4)" below). The lower-dimension vector $\boldsymbol{\delta'}$ was assigned the regularized horseshoe shrinkage prior (annotated as "HS(1)" below) (*63*). We used the "make_stancode" function in the R package brms to generate the Stan code implementing the regularized horseshoe prior. We used the default regularized horseshoe prior implemented by brms (*64*). Our use of a regularized horseshoe prior is justified by the fact that, in the absence of shrinkage on the $\boldsymbol{\delta'}$ prior, many false-positive enrichments would arise. The regularized horseshoe explicitly avoids such problems by applying strong shrinkage of the posterior distribution toward zero, while allowing a small subset of parameter values to explore space further from zero, thus providing a Bayesian analog of "multiple hypothesis testing correction" directly within the structure of the statistical model.

$$\boldsymbol{\alpha}_{g,s} = W_\alpha \cdot \boldsymbol{\alpha}'_{g,s}$$

$$\boldsymbol{\alpha}' \sim \mathrm{N}(0,4)$$

$$\boldsymbol{\delta}_{g,s} = W_\delta \cdot \boldsymbol{\delta}'_{g,s}$$

$$\boldsymbol{\delta}' \sim \mathrm{HS}(1)$$

Finally, the ln(precision) term for each sample type was assigned a normal prior with mean 2 and standard deviation 1.

$$\ln(\phi) \sim \mathrm{N}(2,1)$$

The enrichment scores $\boldsymbol{\beta}$ and input abundance estimates $\boldsymbol{\alpha}$ were converted from the natural-log scale to a $\log_2$ scale prior to interpretation and plotting, and the bedgraph files generated by Enricherator contain these scores on the $\log_2$ scale.

**Running Enricherator**

We ran Enricherator using the R scripts in the Enricherator repository as follows. Note that we have excluded references to file locations from our command line arguments, as these will depend on each user's local computing environment. We direct potential Enricherator users to https://github.com/jwschroeder3/enricherator.git for more detailed usage instructions. We first ran the "cmdstan_fit_enrichment_model.R" script (command line arguments: --ignore_ctgs P2918_rnadna_spikein,NC_011916.1 --norm_method libsize --ext_subsample_dist 30 --ext_fragment_length 60 --input_subsample_dist 60 --input_fragment_length 120 --libsize_key tmm_size_factors). We then ran the script "cmdstan_gather_estimates_from_stanfit.R" (command line arguments: --params Alpha,Beta). We then ran the script "get_contrasts.R" to calculate cross-genotype comparisons (command line arguments: --contrasts rnhB-py97,rnhC-py79,rnhC-rnhB --type genotype). The above commands were run twice, once using strand-specific counts to infer stranded RDH enrichment, and again using non-stranded counts to infer overall RDH enrichment in a non-stranded manner. In many cases we performed additional analyses using robust z-scores of $\log_2$(RDH enrichment) scores. We used our software bgtools (https://github.com/jwschroeder3/bgtools.git) to convert the scores in the bedgraph files output by Enricherator into robust z-scores.

**Statistical modeling of gene expression and orientation's association with RDH enrichment**

For the statistical model used in Figure 2B, we used the R package brms to fit a Bayesian linear model using the formula syntax "*hbd ~ rep*exp*". The model was fit separately for each genotype. The variable *hbd* refers to the gene-wise z-scores for HBD enrichment as determined by Enricherator. The variable *rep* indicates whether a gene is head-on or co-directional, and the variable *exp* indicates the gene-wise z-scores for transcript abundance. The formula syntax above will fit a linear model with the variables, *rep*, *exp*, and and interaction term between *rep* and *exp* as covariates.

**Comparison of RNA:DNA hybrid enrichment in head-on and codirectional genome features**

Genome features are described in the genome annotation file [GenBank accession: NC_022898, (*65*)]. These features were divided into "co-directional" and "head-on" replication vs. transcription orientations based on the origin of replication being at bp 2000 and the replication terminus being at bp 1970043 in the genome coordinates, and the direction of transcription indicated in the annotation file. ncRNA and UTR features were taken from Table S5 of (*66*), filtered for features for which the "Locus_tag" column begins with "BSU_misc" and removing features for which the "classif" column is "-". The work performed in (*66*) was done in the *B. subtilis* 168 strain, so we next mapped the ncRNA and UTR features from *B. subtilis* 168 to the *B. subtilis* PY79 genome using BLAST. We converted the feature records mentioned above to the bed format and retrieved each RNA feature's sequence using "bedtools getfasta". Each resulting fasta sequence was mapped to the *B. subtilis* PY79 reference genome using BLAST, and the BLAST results were converted to bed format, and the assignment of each RNA as ncRNA, 5′-, or 3′-UTR performed by (*66*) was joined to the appropriate features in the resulting bed file, which was finally converted to the gff format. See section SM5.4 above for the url for the github repository containing code used to make the custom annotations.

Feature-level robust z-scores were taken as the median of the position-wise robust z-scores for all positions in that feature. To determine statistical significance of differences occurring between genotypes, two-sided Wilcoxon rank sum tests were performed using the wilcoxon function in the python module scipy.stats. Arguments were zero_method=”wilcox”, correction=False, alternative=”two-sided”, nan_policy=”propagate”. To calculate p-values for comparisons between head-on and codirectional genes within genotypes we used the ranksums function in the python module scipy.stats. Arguments were alternative=”two-sided” and nan_policy=”propagate”. Confidence intervals were calculated via bootstrap sampling. Adjusted p-values (FDR) were generated using the multitest package for R version 3.5.1 (mt.rawp2adjp) using the Benjamini & Hochberg procedures (proc=“BH”).

*SM5.6: qPCR of material used for genome-wide RDH enrichment analysis*

To determine the enrichment of RNA:DNA hybrids via qPCR for wild type (PY79), Δ*rnhB*, and Δ*rnhC* cells, input and extracted material from HBD pull-down were used to assemble qPCR reactions. The reactions were set up to have 1x iTaq Universal SYBR Green Supermix with 300 nmol of each primer set (Data File S1), 0.046875 µL of template per 10 µL reaction. Thermal cycles were 98°C for 5 min, 45 cycles of [ 10 s at 98° C then 30 s at 60°C], followed by a melt curve from 65°C to 95°C in a BioRad CFX Opus RT-PCR instrument. Primers were chosen to span a range of HBD pulldown efficacies and targeted the following areas: upstream regions of *sdpC*, *aroK*, *braB*, *bsrI*, *mfd*, and *cspB*, and the coding region of *ybfM* in *B. subtilis* and the $tRNA^{Thr}$ locus of *C. crescentus*. Cq values were determined via regression using CFX Maestro Software (BioRad).

*SM5.7: Analysis of pPCR RDH enrichment*

We used the python module pymc, version 5.0.2, to infer RHD enrichment vs. input DNA for each set of primers used in section SM5.6. To directly compare ΔΔCq values across genomic loci we interrogated, we devised a statistical model that directly incorporated primer efficiency and pulldown efficiency into the model structure. The data used as inputs to the model and the code can be found at https://github.com/jwschroeder3/qpcr_analysis.git.

*SM6: Bayesian inference of parameter association with mutation rate*

Mutation accumulation data were subset by mutation category, i.e., transition, transversion, or insertion/deletion. A generalized linear model was fit to the data of each mutation category to regress the number of mutations in coding sequences against coding sequence direction relative to the direction of DNA replication fork progression, expression in terms of mean-centered, standardized rpkm values, mean-centered, standardized RNA:DNA hybrid binding domain ChIP enrichment, mean-centered, standardized enrichment of input DNA from our RNA:DNA hybrid binding domain sequencing library preparation, and interaction terms between each of the prior mentioned variables and coding sequence expression. The model fit to the data is summarized as follows:

$$count_i \sim Poisson(\lambda_i)$$

$$\begin{aligned}
\ln(\lambda_i) \quad &= \alpha \\
&+ \ln(l_i \times g) \\
&+ \beta_{HO} \times HO_i \\
&+ \beta_{ex} \times ex_i \\
&+ \beta_{HBD} \times HBD_i \\
&+ \beta_{in} \times in_i \\
&+ \beta_{\Delta rnhB} \times \Delta rnhB_i \\
&+ \beta_{\Delta rnhB:HO} \times \Delta rnhB_i \times HO_i \\
&+ \beta_{\Delta rnhB:ex} \times \Delta rnhB_i \times ex_i \\
&+ \beta_{\Delta rnhB:HBD} \times \Delta rnhB_i \times HBD_i \\
&+ \beta_{\Delta rnhB:in} \times \Delta rnhB_i \times in_i \\
&+ \beta_{\Delta rnhC} \times \Delta rnhC_i \\
&+ \beta_{\Delta rnhC:HO} \times \Delta rnhC_i \times HO_i \\
&+ \beta_{\Delta rnhC:ex} \times \Delta rnhC_i \times ex_i \\
&+ \beta_{\Delta rnhC:HBD} \times \Delta rnhC_i \times HBD_i \\
&+ \beta_{\Delta rnhC:in} \times \Delta rnhC_i \times in_i
\end{aligned}$$

$$\alpha \sim N(-23, 5)$$

$$\begin{bmatrix} \beta_{HO} \\ \beta_{ex} \\ \beta_{HBD} \\ \beta_{in} \\ \beta_{\Delta rnhB} \\ \beta_{\Delta rnhC} \end{bmatrix} \sim N(0,3)$$

$$\begin{bmatrix} \beta_{\Delta rnhB:HO} \\ \beta_{\Delta rnhB:ex} \\ \beta_{\Delta rnhB:HBD} \\ \beta_{\Delta rnhB:in} \\ \beta_{\Delta rnhC:HO} \\ \beta_{\Delta rnhC:ex} \\ \beta_{\Delta rnhC:HBD} \\ \beta_{\Delta rnhC:in} \end{bmatrix} \sim Horseshoe(1)$$

In the above generalized linear model, $count_i$ is the number of mutations observed in CDS i. α is the intercept, and the term $ln\ (l_i \times g)$ serves as an exposure term to account for varying CDS length and the number of generations a given genotype's MA lines underwent. We note that this exposure term causes the fit of the model to our data to infer the association between each covariate and mutation rate per nucleotide per generation, rather than simply mutation count. $HO_i$ is a binary variable taking the value 0 if CDS i is co-directionally transcribed with the direction of DNA replication fork progression and 1 if it is head-on. The variable $ex_i$ represents the expression level of CDS i. To calculate $ex_i$, we calculated $ln\ (rpkm \quad + 1 \times 10^{-5})$ values for each CDS using RNA-seq data from PY79 (wild type), Δ*rnhB*, and Δ*rnhC* strains. Adding $1\text{x}10^{-5}$ to rpkm prior to taking the natural logarithm was done to avoid taking the logarithm of zero for the rare CDSs with no alignments. These $ln\ (rpkm + 1 \times 10^{-5})$ values were then standardized for each genotype's RNA-seq data to determine an $ex$ value for each CDS from each genotype. HBD ChIP enrichment scores were determined as described in Section SM5. For each strain background, HBD pull-down enrichment scores were standardized to determine the value of the variable $HBD$ for each CDS from each genotype.

Input scores were determined as described in section SM5. For each strain background, input scores were standardized to generate $input$ values for each CDS from each genotype. A regularizing prior on α was set as a normal distribution with mean = -23 and standard deviation = 5 to reflect our prior knowledge that the average mutation rate per nucleotide per generation is approximately $e^{-23} \cong 1x10^{-10}$. A normal distribution of mean = 0 and standard deviation = 3 was used as a prior for the main effects of genotype, gene expression z-score, HBD pull-down z-score, input z-score, and gene orientation. Due to the sparse nature of mutation accumulation line data, we used a horseshoe shrinkage prior with degrees of freedom of 1 and scale of 1 for all interaction terms between variables in the model.

We note that because a natural logarithm link function is applied in fitting Poisson generalized linear models such as ours, our point estimate of -22.5 for the intercept of the model fit to transition data (Data File S2) represents an intercept mutation rate of $e^{-22.5}$ per nucleotide per generation, which is approximately $1.69 \times 10^{-10}$ transitions per nucleotide per generation. Additionally, interpretation of the associations between mutation rates and expression, HBD pull-down score, and input score should account for the link function and the fact that the data for these continuous predictor variables are in units standard deviation due to their standardization. To demonstrate this point, let us consider an example where a point estimate for hypothetical variable A's effect on mutation rate is 0.5. This would indicate that if all other variables were maintained at a constant value, a one-standard-deviation increase in the value of variable A would correspond to a relative increase in average predicted mutation rate by a factor of $e^{0.5}$.

Inference of each variable's association with CDS mutation rate was performed using the R package "brms" (*64*), which generates code in the programming language Stan to fit a model using Bayesian inference (*61*). The data used to fit the model are in Data File S2. Posterior samples for each variable were extracted from the resulting fit objects and summarized using the R package "tidybayes". Point estimates (median values for a given posterior distribution), 90% highest continuous probability intervals, and evidence ratios (Bayes factors, or K) for all parameters' associations with mutation rates and hypotheses tested are in Data File S2.

Hypothesis testing was performed by using the function "hypothesis" from the R package "brms" (*64*). We note that the Bayesian interpretation here provides a measure of support for a hypothesis of interest, and not simply against a specified null hypothesis. We therefore tested three hypotheses for each parameter's estimated association with mutagenesis: 1) the parameter is positively associated with mutation rates ("Evidence ratio positive" column in Data File S2), 2) the parameter is negatively associated with mutation rates ("Evidence ratio negative" column in Data File S2), and 3) the parameter's association with mutation rates is close to zero ("Evidence ratio zero" column in Data File S2). For the hypothesis that the association is near zero, we defined a region of practical equivalence from (-ln(1.2),ln(1.2)). For the directed hypotheses, significance symbols took the form of "*" symbols. For the hypothesis that the association is near zero, we use "X" symbols. The evidence ratios (K) were calculated as the proportion of values from a posterior distribution that met the hypothesis divided by the proportion of values from the posterior distribution that violated the hypothesis. We applied Kass and Raftery's scale for defining strength of evidence, by which a K-value from [3,20) indicates "positive" evidence in support of the hypothesis and the parameter receives a single significance symbol (* or X), a K-value from [20,150) represents "strong" evidence and the parameter receives two significance symbols (** or XX), and a K-value from [150,Inf) indicates "very strong" evidence in support of the hypothesis and the parameter receives three significance symbols (*** or XXX) (*33*). These ranges can be interpreted as providing similar value to particular p-value ranges that might arise in frequentist statistical testing. For example, the K=3 cutoff in the Bayesian framework is analogous to the commonly used p=0.05 cutoff in frequentist tests.

In addition to the model described above (we call this the “full model”), we fit an additional four modified versions of the model. 1) A model without any term referencing gene expression, 2) a model without any term referencing RDH enrichment, 3) a model with two RDH slopes for the main effect of RDH enrichment and each interaction term for which RDH enrichment was a member, the changepoint between the two slopes was also an additional fitted parameter, 4) a model with a slope of zero to the left of a fitted changepoint, and a fitted slope associated with RDH enrichment to the right of the changepoint. We found that for all three mutation types analyzed in our work, the full model described above, without any changepoint of RDH association with mutation rate, was either the best-performing model or was within the standard error of the best performing model as judged using the “loo_compare” function in version 2.5.1 of the “loo” package **(Table S1)**. We therefore saw no evidence-based reason to use any of these four additional models, and instead used the full model defined above.

### Supplementary Figures

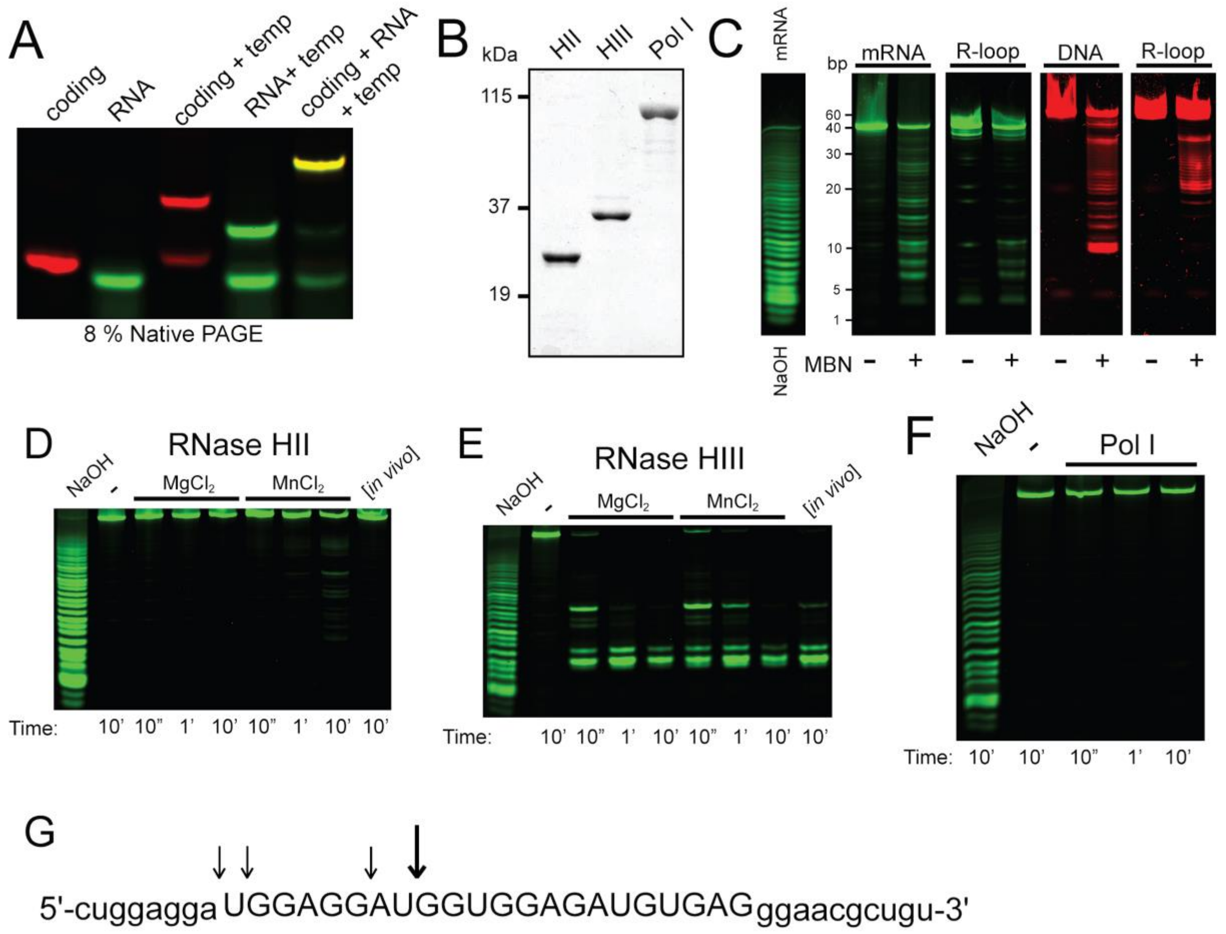


**Figure S1 (related to Figure 1) Assembly of the R-loop substrate. (A)** Native PAGE of different combinations of the oligonucleotides showing sequential R-loop assembly. **(B)** Purity of recombinant RNase HII (RnhB), RNase HIII (RnhC), and pol I (PolA) shown via SDS-PAGE using 2 mg total protein. **(C)** Urea-PAGE of Mung Bean Nuclease (MBN), a single stranded RNA and DNA endo- and exonuclease, digestion of the mRNA strand (green) and template strand (red) either with or without the full R-loop complex. NaOH-induced hydrolysis of the substrate RNA oligonucleotide. **(D-E)** Urea-PAGE of the R-loop incubated with (D) 4 nM RnhB (HII) or (E) 4 nM RnhC in the presence of 1 mM $MgCl_2$, 1 mM $MnCl_2$ or both metals present at physiologically relevant concentrations (1 mM $MgCl_2$ and 10 µM $MnCl_2$ lane labeled *in vivo*). **(F)** Urea-PAGE of the R-loop incubated with 4 nM pol I over time. This gel indicates that the R-loop is stable and maintained over a 10 min time-course despite the introduction of a protein purified using the same procedure as RnhB (HII) and RnhC (HIII). **(G)** Arrows showing the locations of RnhC-mediated hydrolysis of the mRNA sequence. Arrow length denotes likelihood of cleavage at that site.

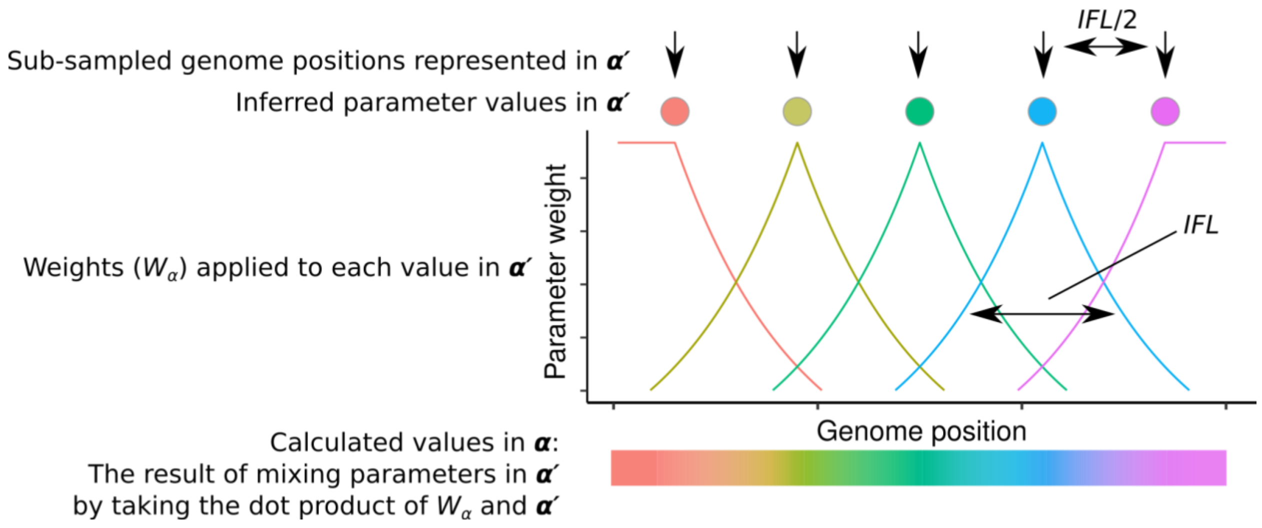


**Figure S2 (related to Figure 2). Graphical depiction of parameter smoothing in Enricherator.** *IFL* = mean input fragment length; $\boldsymbol{\alpha}$ = vector of inferred values for input DNA coverage, calculated using $\boldsymbol{\alpha'}$ and $W_{\alpha}$; $\boldsymbol{\alpha'}$ = lower-dimensional vector of fitted values at resolution *IFL*/2. Filled circles represent the parameter value for each element of the vector $\boldsymbol{\alpha'}$, with each distinct color representing a distinct fitted value. The color of each exponential kernel indicates the element of $\boldsymbol{\alpha'}$ to which the given kernel applies when taking the dot product, $\boldsymbol{\alpha} = W_{\alpha} \cdot \boldsymbol{\alpha'}$. For HBD pull-down data the logic is identical; the fragment length (*EFL*) may be different than the input fragment length.

A

| Strain | cells scored | % RecA-GFP | cells scored | % TagC-GFP |
|---|---|---|---|---|
| Wild type | 1589 | 10.6 ± 4.3 | 1839 | 0.63 ± 0.23 |
| Δ*rnhC* | 1050 | 26.4 ± 6.3 | 1718 | 12.0 ± 6.9 |

B

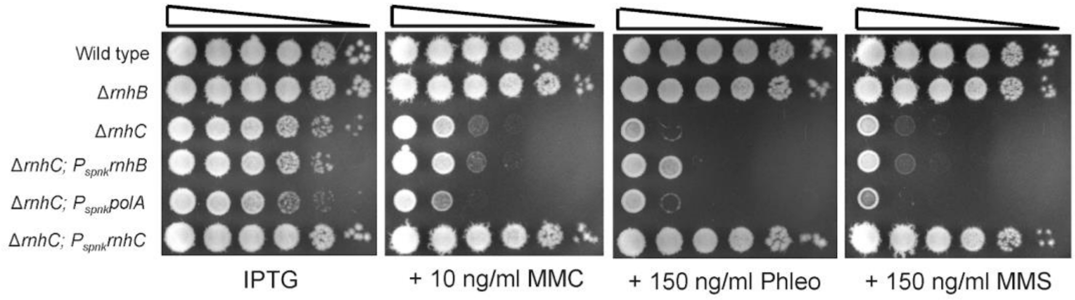


**Figure S3 (related to Figure 3). Cells lacking *rnhC* are sensitive to DNA damage and SOS induced. (A)** RecA-GFP and TagG-GFP fluorescence data of wild type and Δ*rnhC* cells during normal growth in the absence of DNA damage. **(B)** Spot titer assays of various RNase H deficient strains serially diluted to $10^{-5}$ and plated on 1 mM IPTG (left) and IPTG plus MMC, phleomycin, or MMS.

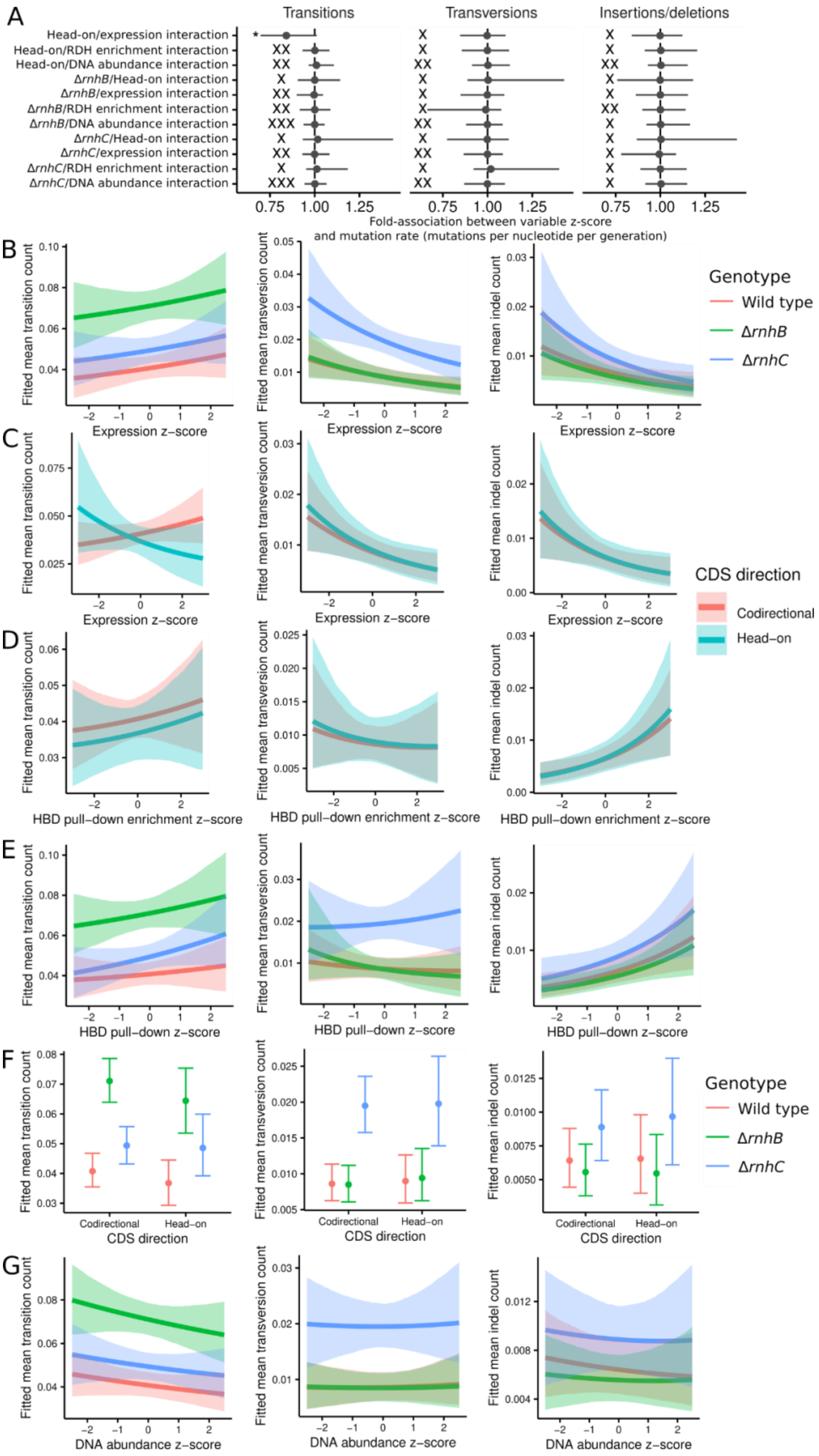
A
Transitions
Transversions
Insertions/deletions
Head-on/expression interaction
Head-on/RDH enrichment interaction
Head-on/DNA abundance interaction
ΔrnhB/Head-on interaction
ΔrnhB/expression interaction
ΔrnhB/RDH enrichment interaction
ΔrnhB/DNA abundance interaction
ΔrnhC/Head-on interaction
ΔrnhC/expression interaction
ΔrnhC/RDH enrichment interaction
ΔrnhC/DNA abundance interaction
Fold-association between variable z-score
and mutation rate (mutations per nucleotide per generation)
B
C
D
E
F
G
Fitted mean transition count
Fitted mean transversion count
Fitted mean indel count
Expression z-score
HBD pull-down enrichment z-score
HBD pull-down z-score
CDS direction
DNA abundance z-score
Codirectional
Head-on
Genotype
Wild type
ΔrnhB
ΔrnhC
CDS direction
Codirectional
Head-on

**Figure S4 (related to Figure 4) Analysis of genome-wide mutation rates. (A)** Inferred associations between the rate of transition, transversion, or indel occurrence in CDSs and each of our statistical model's interaction terms. Main effects are in Figure 4. Stars indicate the strength of evidence for a parameter's association with the natural log of mutation rate being either greater or less than zero. An "X" indicates the strength of evidence for a parameter's association with the natural log of mutation rate being near zero. Strength of evidence was determined as described in Materials and Methods. The symbol * represents "positive" support with a K-value [3,20), ** represents a K-value from [20,150) indicating "strong" evidence for the parameter, and *** represents a K-value from [150,Inf) indicating "very strong" evidence for a hypothesis; equivalently strong $K_0$ values are shown with equal numbers of 'X' characters. **(B-G)** Fitted values for mean transition, transversion, or indel count for all marginal effects for a theoretical average CDS implied by our model. Points or lines represent the median estimate and bars or shaded intervals denote the 90% highest-density continuous interval.

| **Model name** | **Mutation type** | **LOO-IC (SE)[a]** | **ELPD difference (SE)[b]** |
|---|---|---|---|
| **Full model** | **transitions** | **4821 (146.1)** | **-0.5462 (1.143)** |
| **No RDH** | **transitions** | **4820 (146.1)** | **0.0 (0.0)** |
| **No expression** | **transitions** | **4822 (146.2)** | **-0.7385 (2.286)** |
| **Two RDH slopes** | **transitions** | **4824 (146.2)** | **-1.934 (1.269)** |
| **Right RDH slope** | **transitions** | **4852 (147.7)** | **-15.84 (6.237)** |
| **Full model** | **transversions** | **1573 (109.4)** | **-0.4593 (0.6261)** |
| **No RDH** | **transversions** | **1572 (109.3)** | **0.0 (0.0)** |
| **No expression** | **transversions** | **1575 (109.9)** | **-1.820 (2.823)** |
| **Two RDH slopes** | **transversions** | **1577 (109.9)** | **-2.383 (0.8837)** |
| **Right RDH slope** | **transversions** | **1574 (109.4)** | **-0.9705 (0.5135)** |
| **Full model** | **indels** | **1060 (99.05)** | **0.0 (0.0)** |
| **No RDH** | **indels** | **1065 (99.32)** | **-2.570 (2.524)** |
| **No expression** | **indels** | **1064 (99.02)** | **-1.913 (2.437)** |
| **Two RDH slopes** | **indels** | **1069 (100.0)** | **-4.509 (2.665)** |
| **Right RDH slope** | **indels** | **1061 (99.18)** | **-0.3868 (0.8443)** |

**Table S1.** Results of quantitative model comparison for the fits of the five indicated models to each of the three indicated mutation rates for mutations of the indicated type. Values were calculated using version 2.5.2 of the "loo" package in R. **a)** LOO-IC = LOO information criterion. Lower values of LOO-IC indicate better predictive performance. **b)** ELPD difference = difference between expected log pointwise predictive density for the model with the best predictive performance versus the given model/mutation type. A value of 0.0 is associated with the model with the best predictive performance, and increasingly negative values indicate increasingly poorly performing models. Note that across mutation types the "full model" is either the best-performing model, or the standard error for the ELPD difference for the "full model" always overlaps the ELPD difference for the best model, indicating their performances are indistinguishable.

**Data File S1.** Excel workbook containing multiple sheets. Contains strains, oligonucleotides, and qPCR primers used in this study.

**Data File S2.** Excel workbook containing multiple sheets. Contains an overview of the mutations identified in MA lines, details of the structural variants identified in MA lines, the data used to fit our statistical models for CDS mutation rate, fitted parameter estimates from our statistical model, and results of RNA-seq analysis for *rnhC::erm, lexA[G92D]* vs. wild type, Δ*lexA* vs. wild type, Δ*rnhC* vs. wild type, and Δ*rnhB* vs. wild type cells.
